# Comparative plant transcriptome profiling of Arabidopsis and Camelina infested with *Myzus persicae* aphids acquiring circulative and non-circulative viruses reveals virus- and plant-specific alterations relevant to aphid feeding behavior and transmission

**DOI:** 10.1101/2022.01.04.474350

**Authors:** Quentin Chesnais, Victor Golyaev, Amandine Velt, Camille Rustenholz, Véronique Brault, Mikhail Pooggin, Martin Drucker

## Abstract

**Background:** Evidence accumulates that plant viruses alter host-plant traits in ways that modify their insect vectors’ behavior. These alterations often enhance virus transmission, which has led to the hypothesis that these effects are manipulations caused by viral adaptation. However, the genetic basis of these indirect, plant-mediated effects on vectors and their dependence on the plant host and the mode of virus transmission is hardly known.

**Results:** Transcriptome profiling of *Arabidopsis thaliana* and *Camelina sativa* plants infected with turnip yellows virus (TuYV) or cauliflower mosaic virus (CaMV) and infested with the common aphid vector *Myzus persicae* revealed strong virus- and host-specific differences in the gene expression patterns. CaMV infection caused more severe effects on the phenotype of both plant hosts than did TuYV infection, and the severity of symptoms correlated strongly with the proportion of differentially expressed genes, especially photosynthesis genes. Accordingly, CaMV infection modified aphid behavior and fecundity stronger than did infection with TuYV.

**Conclusions:** Overall, infection with CaMV – relying on the non-circulative transmission mode – tends to have effects on metabolic pathways with strong potential implications for insect-vector / plant-host interactions (e.g. photosynthesis, jasmonic acid, ethylene and glucosinolate biosynthetic processes), while TuYV – using the circulative transmission mode – alters these pathways only weakly. These virus-induced deregulations of genes that are related to plant physiology and defense responses might impact aphid probing and feeding behavior on both infected host plants, with potentially distinct effects on virus transmission.

## Introduction

Most plant viruses rely on vectors for transmission to a new host (for example Dietzgen et al., 2016). Phloem-feeding insects, such as whiteflies and aphids, are important vectors transmitting at least 500 virus species (Fereres and Raccah, 2015). The high virus transmission capacity is due to their particular non-destructive feeding behavior that allows virus acquisition from and inoculation into the cytoplasm and/or the phloem sap of a new host plant. In fact, aphids alighting on a new plant will first evaluate the potential host for suitability by exploratory intracellular punctures into the epidermis and underlying tissues. If the plant is accepted, aphids plunge their needle-like mouthparts, the so-called stylets, for prolonged feeding phases into the sieve cells whose sap constitutes their principal nutritive source (for review Dáder et al., 2017). Aphids secrete different saliva types during both the probing and the feeding phases that contain effector molecules controlling interactions with the plant and susceptibility (Rodriguez and Bos, 2013).

Viruses are classified according to two principal modes of transmission (for review Blanc et al., 2014). Circulative viruses such as turnip yellows virus (TuYV, genus *Polerovirus*) are acquired by aphid (or other insect) vectors from the phloem sap of infected plants. They bind to specific receptors on the intestine epithelium (Mulot et al., 2018), traverse the intestine and cycle through the hemocoel to subsequently reach and invade the salivary glands (Brault et al., 2007). New hosts are inoculated when viruliferous aphids migrate between plants and inoculate the virus during salivation phases into the phloem, the only tissue where TuYV and many other circulative viruses are able to replicate. For this mode of transmission, virus acquisition and inoculation periods are rather long (in the range of several hours to days), requiring that aphids settle sustainably on the plants. On the other hand, despite the fact that poleroviruses do not seem to replicate in the vector, aphids having acquired poleroviruses remain infectious during their entire lifespan. Therefore, this transmission mode is also referred to as persistent transmission.

The transmission mode of non-circulative viruses such as cucumber mosaic virus (CMV, genus *Cucumovirus*), that are also transmitted by aphids and other hemipteran vectors, is entirely different (for review Ng and Falk, 2006)). They are mostly acquired and inoculated during early probing phases [i.e. intracellular penetrations in the epidermis and mesophyll tissues (Martin et al., 1997)]. They do not invade aphid cells but are retained externally in the mouthparts (stylets and/or esophagus), from where they are also released into a new host. For this reason, non-circulative viruses are acquired and inoculated within seconds to minutes, and vectors retain and transmit the virus only for a limited time (minutes range). Therefore, this transmission mode is also named non-persistent transmission. Some non-circulative viruses may be retained by the vectors during several hours and are referred to as semi-persistent viruses. The aphid-transmitted cauliflower mosaic virus (CaMV, genus *Caulimovirus*) belongs to this group (Kennedy et al., 1962; Moreno et al., 2012).

Available data indicate that many viruses modify host traits (i.e. color, volatiles, primary/secondary metabolites etc.) in ways that are conducive for their transmission (for review Dáder et al., 2017; Fereres and Moreno, 2009; Mauck et al., 2012). Theoretical considerations suggest that these modifications depend on the virus species and in particular on the mode of virus transmission by vectors (Dáder et al., 2017; Mauck et al., 2012). Circulative, persistent and phloem-restricted viruses should profit in particular from faster access of vectors to the phloem and longer feeding, which would promote both virus acquisition and inoculation. In addition, these viruses would tend to improve nutrient quality of the host and consequently vector fitness and fecundity, concomitant with an increased number of viruliferous vectors (Dáder et al., 2017; Fereres and Moreno, 2009; Mauck et al., 2018). Both modifications have been reported for aphid-transmitted luteoviruses (Bosque-Pérez and Eigenbrode, 2011). Non-persistent or semi-persistent, non-circulative viruses with their fast transmission kinetics are expected to benefit from the attraction of vectors to infected plants, followed by a rapid dispersion, before the virus is lost from the vector during subsequent salivation events. The best-studied example is CMV, where altered volatiles incite aphids to alight on infected plants and acquire CMV, before the poor taste and low nutritive value encourage the aphids to leave and transmit the virus, attached to the stylets during this brief probing time, to other (healthy) host plants (Carr et al., 2020; Mauck et al., 2010; Mauck et al., 2014).

While there is overwhelming evidence that some viruses do achieve plant phenotype manipulation in ways that are conducive for their own transmission, many significant knowledge gaps remain (discussed by Mauck and Chesnais, 2020). In particular, the mechanisms and pathways by which viruses alter aspects of the host phenotype and the virus components that manipulate the host remain poorly understood (Mauck et al., 2019).

In the present study, we addressed these shortcomings and initiated analysis of the effects of two viruses, TuYV and CaMV, belonging to two different transmission categories, on transcriptomic profiles in two susceptible host plant species (*Arabidopsis thaliana* and *Camelina sativa*, both family *Brassicaceae*), and on changes in insect vector feeding behavior and performances. We selected the green peach aphid *Myzus persicae* as vector because it transmits both TuYV and CaMV and infests both plant hosts. We chose two different plant species as virus hosts, as previous studies have highlighted potential host-specific effects of viruses on host plant traits and vector performance (Chesnais et al., 2019b; Chesnais et al., 2021). In addition, their phylogenetic proximity allows rather easy gene-to-gene comparisons. In fact, the *C. sativa* genome is highly similar to the *A. thaliana* genome and might have arisen from hybridization of three diploid ancestors of *A. thaliana* (Malik et al., 2018). For this reason, its genome is allohexaploid. Over 70 % of *C. sativa* genes are syntenically orthologous to *A. thaliana* genes (Kagale et al., 2014), facilitating genomic studies of *C. sativa*.

## Material and methods

### Aphids

A Dutch green peach aphid clone (*Myzus persicae* Sulzer, 1776) was used for the experiments. It was reared on Chinese cabbage (*Brassica rapa* L. *pekinensis* var. Granaat) in a growth chamber at 20±1 °C and a 16 h photoperiod. Only wingless forms were used in assays. For synchronization, adults were placed on detached Chinese cabbage leaves that were laid on 1 % agarose in a Petri dish. The adults were removed 24 h later and the newborn larvae used in experiments 5 days (for transcriptomic experiments) or 8 days (for feeding behavior and performances experiments) later.

### Viruses

CaMV isolate Cm1841-Rev (Chesnais et al., 2021), which is a transmissible derivative of isolate Cm1841 (Tsuge et al., 1994), and TuYV isolate TuYV-FL1 (Veidt et al., 1988) were maintained in *A. thaliana* Col-0 and propagated by aphid inoculation of 2-week-old plants. Growth conditions were as described below.

### Virus infection and aphid infestation

Seeds of *Arabidopsis thaliana* Col-0 (hereafter Arabidopsis) or *Camelina sativa* var. Celine (hereafter Camelina) were germinated in TS 3 fine substrate (Klasmann-Deilmann) in 7*7 cm pots and watered with tap water. Growth conditions were 14 h day 10 h night with LED illumination and a constant temperature of 21±1 °C. Two-week-old plants were inoculated with 3-5 wingless *Myzus persicae* aphids that had been allowed a 24 h acquisition access period on Arabidopsis infected with TuYV or CaMV or on healthy Arabidopsis. Plants were individually wrapped in clear plastic vented bread bags to prevent cross contamination. Aphids were manually removed after a 48 h inoculation period. Eighteen days post-inoculation (dpi), 25 to 30 5-day-old non-viruliferous aphids were placed for infestation on the rosette (Arabidopsis) or the apical leaves (Camelina) of CaMV-or TuYV-infected or mock-inoculated plants. After 72 h infestation (= 21 dpi), aphids were removed with a brush. The infested plants (virus-infected or mock-inoculated) were washed 3 times with deionized water and 3 times with MilliQ water to remove any remaining aphid exuvia and honeydew. Then rosettes (Arabidopsis) or detached leaves (Camelina) were collected in 50 ml Falcon tubes. Three biological replicates were used for analysis. For Arabidopsis, one biological replicate consisted of 4 plants, for Camelina one replicate was 3 plants. Plant samples were conserved at −80 °C until processing.

### RNA purification and Illumina sequencing

Total RNA was extracted from one gram of Arabidopsis (rosettes) and Camelina (leaves) frozen tissues using a CTAB-LiCl protocol (Morante-Carriel et al., 2014) modified as described in detail by Golyaev et al. (2019). Briefly, the plant material was ground in liquid nitrogen, homogenized in 10 ml CTAB buffer and centrifuged for 10 min at 5,000 g and 4 °C. The supernatant was mixed with one volume of chloroform:isoamyl alcohol (24:1) followed by nucleic acid precipitation with 0.1 volume of 3 M sodium acetate (NaOAc, pH 5.2) and 0.6 volume of isopropanol, incubation at −20 °C for 1 h and centrifugation for 20 min at 20,000 g and 4 °C. The pellet was resuspended in 1 ml of RNase-free water followed by selective precipitation of RNA by addition of 0.3 volume of 10 M LiCl, overnight incubation at 4 °C and centrifugation for 30 min at 20,000 g and 4 °C. The RNA pellet was resuspended in 0.1 ml of RNase-free water, 0.1 volume of 3 M NaOAc (pH 5.2) and 2 volumes of cold absolute ethanol, centrifuged for 20 min at 20,000 g and 4 °C, washed with ice-cold 70 % ethanol, air-dried and dissolved in 50 μl RNAse-free water.

Total RNA samples were subjected to quality control and Illumina sequencing at Fasteris (www.fasteris.com) using a standard, stranded mRNA library preparation protocol and multiplexing the resulting 18 libraries (3 biological replicates per each of the six conditions) in one NovaSeq flowcell SP-200 with 2×75 nt paired-end customized run mode. The resulting 75 nt reads from each library were mapped with or without mismatches to the reference genomes of Arabidopsis (TAIR10.1 nuclear genome (5 chromosomes), chloroplast (Pltd) and mitochondrion (NT): https://www.ncbi.nlm.nih.gov/genome/?term=txid3702[Organism:noexp]), Camelina (nuclear genome (20 chromosomes): https://www.ncbi.nlm.nih.gov/genome/?term=txid71323[Organism:exp]), CaMV [strain CM1841rev (Chesnais et al., 2021)] and TuYV [NC_003743, (Veidt et al., 1988)]. Note that the CaMV reference sequence was extended at the 3’-end by 74 nts from the 5’-terminus to account for its circular genome and allow for mapping reads containing the first and last nucleotides of the linear sequence. In the case of TuYV, some discrepancies with the reference sequence were detected, when the reads were mapped to the viral reference sequence. Therefore, the reads were used to generate a new consensus master genome in the viral quasispecies population. For both viruses, the consensus genome sequences (Supplementary Sequence information S1) were used for (re-)mapping and counting total viral reads as well as viral reads representing forward and reverse strands of the viral genomes (Supplementary Dataset S1).

### RT-qPCR

Expression of Arabidopsis genes was monitored by RT-qPCR analysis. cDNA was synthesized from 10 μg total RNA using AMV Reverse Transcriptase (Promega) and oligo-dT. Real-time qPCR reactions were completed in the LightCycler^®^ 480 instrument (Roche) using the SybrGreen master mix (Roche) and following the recommended protocol. Each reaction (10 μl) included 3 μl of cDNA and 0.5 μl of 10 μM primers (Supplementary Table S1). The thermocycler conditions were as follows: pre-incubation at 95 °C for 5 min, followed by 40 cycles of 95 °C for 10 s, 58-60 °C for 20 s and 72 °C for 20 s. The expression was normalized to the Arabidopsis internal reference gene PEX4 (AT5G25760) (Supplementary Table S1).

### Raw data processing and quality control for transcriptome profiling

Processing was carried out on the Galaxy France platform (https://usegalaxy.fr/) (Afgan et al., 2016). Raw reads quality were checked with FastQC (v0.11.8) and the results were then aggregated with MultiQC (v1.9). For Arabidopsis, between 58.6 and 69.4 million 75 nt paired-end reads were sequenced with a mean phred score >30 for all bases. For Camelina, between 56.4 and 77.6 million 75 nt paired-end reads were sequenced with a mean phred score >30 for all bases. In all samples, there were no overrepresented sequences and really few adapter (0.15% of adapter for the last bases of reads). Reads are aligned on the reference genome with STAR (v2.7.6a) using default parameters and quality again checked with MultiQC. Between 80% and 92.3% of reads were uniquely mapped for Arabidopsis samples and between 60.8% and 70.5% of reads were uniquely mapped for Camelina. Between 17 to 20% of reads mapped to multiple loci in Camelina because of the triplication event of this genome. Reference genomes were Camelina_sativa.Cs.dna.release-49 and Arabidopsis_thaliana.TAIR10.49 from EnsemblPlant portal. Gene counts were obtained with featureCounts (v2.0.1). This option allows reads to map to multiple features for Camelina). 92.2% to 93.3% of uniquely aligned reads were assigned to a gene for Arabidopsis and 80.7% to 88.6% aligned reads were assigned to a gene for Camelina. Differential gene expression was then analyzed with SARTools (v1.7.3) and the DESeq2 method (i.e., TuYV-infected plants vs. mock-inoculated, CaMV-infected plants vs. mock-inoculated). GO enrichment analysis was performed with GOseq (v1.36.0) on the DEGs.

To measure viral RNA loads in plants, the RNA-seq reads from each sample were mapped to the reference genome sequences of the host plant (Arabidopsis or Camelina) and the virus (CaMV or TuYV) with zero mismatches, and the mapped reads were sorted by polarity (forward, reverse and total) and counted. Viral read counts were then normalized per million of total (viral + non-viral) or plant reads (see Supplementary Dataset S1).

### Aphid feeding behavior

To investigate the effects of TuYV and CaMV plant infections on the feeding behavior of *M. persicae*, we used the electrical penetration graph technique (EPG) (Tjallingii, 1988). Eight adult aphids were connected to a Giga-8 DC-system (EPG Systems, www.epgsystems.eu) and placed on the leaf of an individual experimental host plant. To create electrical circuits that each included a plant and an aphid, we tethered each insect by attaching a 12.5 μm diameter gold wire to the pronotum using conductive water-based silver glue. The whole system was set up inside a Faraday cage located in a climate-controlled room held at 21±1 °C and under constant LED illumination during recording. Plants were obtained as described in the previous section but, unlike the plants used for the RNA-seq experiment, the plants used in EPG were not pre-infested. We used the PROBE 3.5 software (EPG systems, www.epgsystems.eu) to acquire and analyze the EPG waveforms as described (Tjallingii and Hogen Esch, 1993). Relevant EPG variables were calculated with EPG-Calc 6.1 software (Giordanengo, 2014). We chose variables based on five different EPG waveforms corresponding to: “probing duration”, “stylet pathway phase”, “phloem sap ingestion”, “time to first phloem sap ingestion” and “salivation in phloem sap”. For each aphid x plant x virus combination, we collected 8-hour recordings from 20 to 23 individuals.

### Aphid fecundity

To investigate the effects of TuYV and CaMV plant infections on the fecundity of *M. persicae*, we randomly selected synchronized wingless adults (8±1 day-old) and transferred them onto experimental host plants. For Arabidopsis experiments, we used one plant per aphid, and we covered the pots with vented bread bags. For Camelina experiments, to force aphids to settle on symptomatic leaves, we placed adults on detached leaves that were laid on 1 % agarose in a Petri dish. The number of nymphs produced per adult was recorded after 5 days. We discarded from the analysis the adult aphids that died before the end of the experiment. Data on both Arabidopsis and Camelina host-plants were collected in three repetitions, comprising altogether 27-33 aphids per aphid x plant x virus combination.

### Statistical analyses of aphid behavior and fecundity

Data on aphid feeding behavior were analyzed using generalized linear models (GLMs) with a likelihood ratio and the chi-square (χ^2^) test. Since duration parameters (i.e. probing duration, stylet pathway phase, phloem sap ingestion and salivation) were not normally distributed, we carried out GLMs using a gamma (link = “inverse”) distribution. For the “time to first phloem phase”, we used the cox proportional hazards model and we treated cases where the given event did not occur as censored. The assumption of validity of proportional hazards was checked using the functions “coxph” and “cox.zph”, respectively (R package “survival”). For aphid fecundity, count data were not normally distributed. Accordingly, we carried out a GLM using a Poisson distribution, a quasi-likelihood function was used to correct for overdispersion, and Log was specified as the link function in the model. When a significant effect of one of the main factors was detected or when an interaction between factors was significant, a pairwise comparison using estimated marginal means (R package “emmeans”) (p value adjustment with Tukey method) at the 0.05 significance level was used to test for differences between treatments. The fit of all GLMs was controlled by inspecting residuals and QQ plots. All statistical analyses were performed using R software v. 4.0.4 (www.r-project.org/).

## Results and discussion

### Plant phenotype

We used in this study 5-week-old Arabidopsis or Camelina plants that had been inoculated with CaMV or TuYV 3 weeks before analysis. In both Arabidopsis and Camelina plants, CaMV caused severe leaf curling, mosaic and vein chlorosis as well as dwarfism (Figure 1). TuYV-infected Arabidopsis and Camelina plants were smaller compared to mock-inoculated plants, but showed no leaf deformation or bleaching. Older TuYV-infected Arabidopsis leaves turned purple, probably due to stress-induced anthocyanin accumulation as previously reported for infection of Arabidopsis with another polerovirus, brassica yellows virus (Chen et al., 2018). The purple coloring was first visible on the abaxial leaf surface and progressed slowly until covering the entire leaf very late in infection. Old leaves of TuYV-infected Camelina displayed mild yellowing symptoms, primarily on the leaf border.

**Figure 1:**
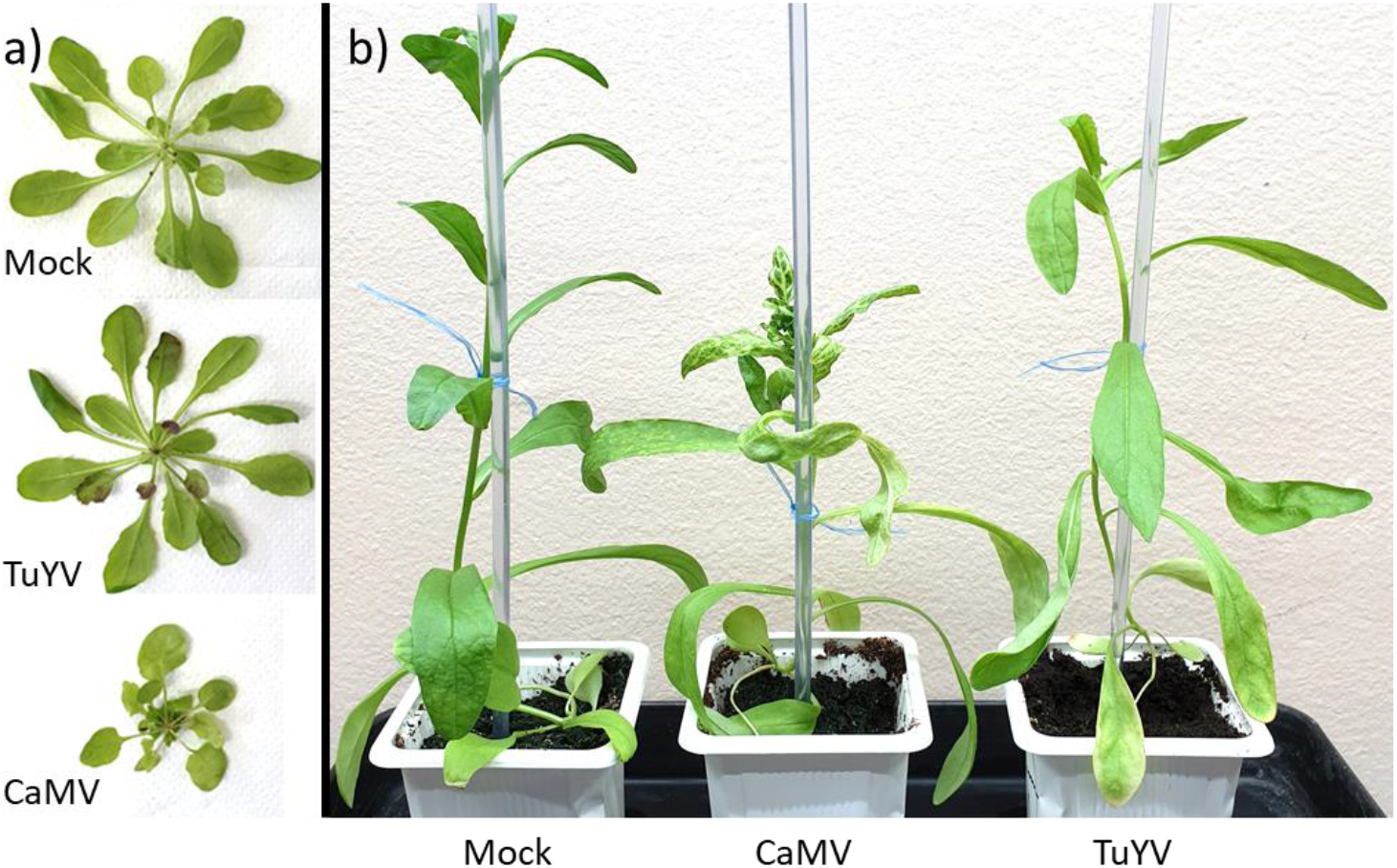
Phenotype of CaMV and TuYV-infected plants. a) Arabidopsis and b) Camelina plants at 21 days after inoculation with the indicated virus or after mock inoculation.

### Aphid feeding behavior and fecundity

We used EPG to compare aphid probing and feeding behavior on Arabidopsis and Camelina infected or not with CaMV or TuYV (Figure 2a,b). The total probing time was identical for all six conditions and the aphids were active for approximately seven hours during the eight hours observation period. The pathway phase and the time until the first phloem ingestion were in general longer on Camelina for all three conditions, whereas the ingestion phase was longer on Arabidopsis. The most important difference was the salivation time, which was extended on Camelina (50-100 % longer than on Arabidopsis), independently of the infection status.

**Figure 2:**
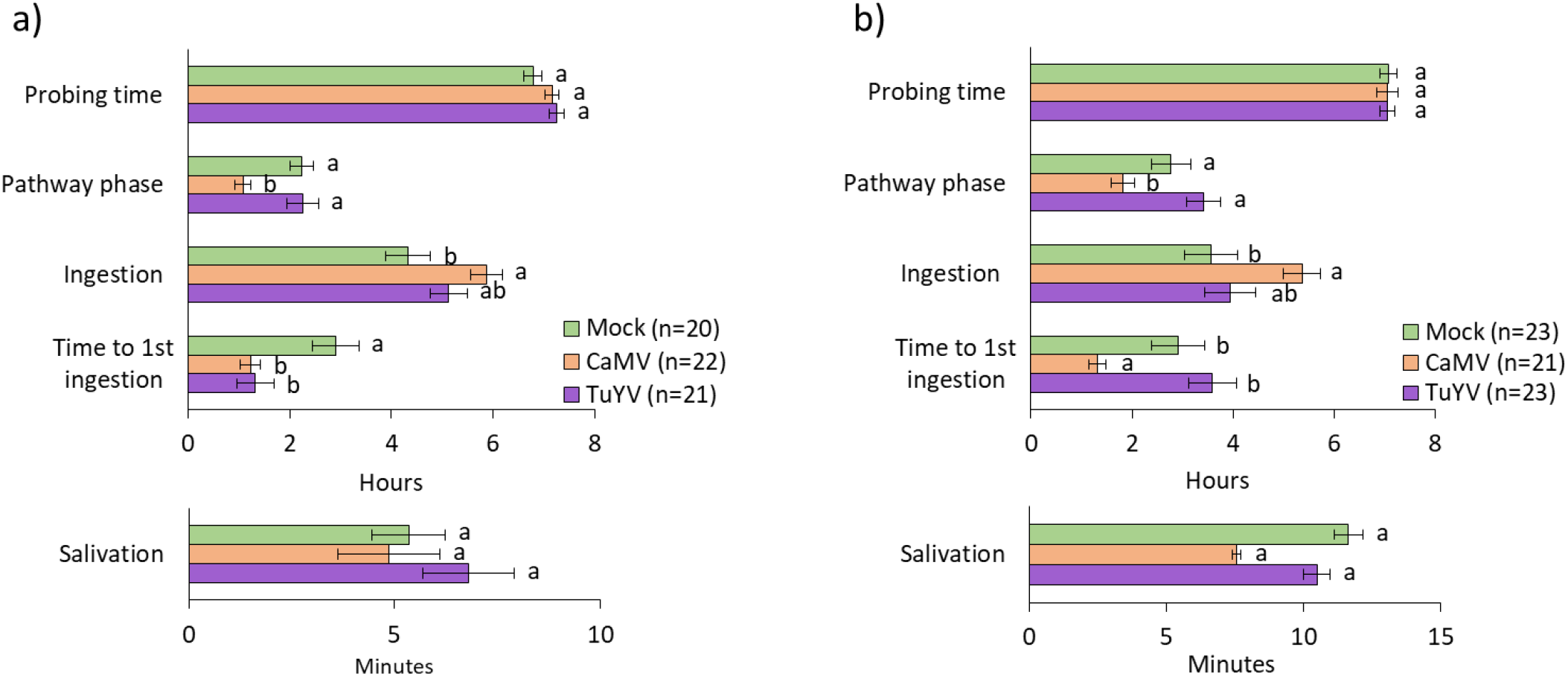
Aphid feeding behavior parameters recorded by EPG on 5-week-old mock-inoculated, TuYV- or CaMV-infected a) Arabidopsis and b) Camelina. Different letters indicate significant differences between plants as tested by GLM followed by pairwise comparisons using “emmeans” (p < 0.05; method: Tukey, n = 20-23).

CaMV infection changed aphid behavior similarly on both plant hosts. The pathway phase and the time to first phloem ingestion were decreased, whereas phloem ingestion was increased. Salivation time was not affected by the infection status of the two-plant species. Infection with TuYV had no major effect. The only significantly affected behavioral parameter was the time until first phloem ingestion, which was reduced by half on TuYV-infected Arabidopsis but not on TuYV-infected Camelina, compared to mock-inoculated plants. This is in contrast with CaMV infection for which the time to first phloem sap ingestion was reduced on both hosts. Previous EPG experiments on Arabidopsis (Bogaert et al., 2020) and Camelina (Chesnais et al., 2019a) have reported neutral to slightly positive effects of TuYV infection on aphid probing and feeding behavior, and highlighted also host-specific viral effects on plant quality and vector behavior (Chesnais et al., 2019b).

Taken together, the significantly reduced time until first phloem ingestion observed on infected Arabidopsis might contribute to a better acquisition of CaMV and TuYV. The other transmission-related feeding parameters were only marginally modified on TuYV-infected plants, whereas CaMV infection altered aphid feeding more strongly. The reduced pathway phase and the increased phloem ingestion might also facilitate CaMV acquisition from phloem tissues. These alterations are expected to be detrimental for non-circulative viruses (such as the non-persistent potyviruses) that are acquired during intracellular penetrations occurring in the pathway phase, but lost if the aphid stylets reach the phloem sap (Kloth and Kormelink, 2020). However, this does not apply to CaMV, acquired efficiently from phloem sap as well as mesophyll and epidermis cells (Palacios et al., 2002).

Infection with CaMV reduced aphid fecundity significantly on both plant host species (Figure 3a,b), compared to mock-inoculated plants (GLM, χ^2^ = 0.0007 and χ^2^ = 0.0409 for Arabidopsis and Camelina, respectively) and correlated with the strong symptoms of infected plants. Fecundity was unchanged on TuYV-infected Arabidopsis and Camelina. This indicated that the severe (but less strong, compared to CaMV infection) phenotype of TuYV-infected Camelina did not interfere with aphid fecundity.

**Figure 3:**
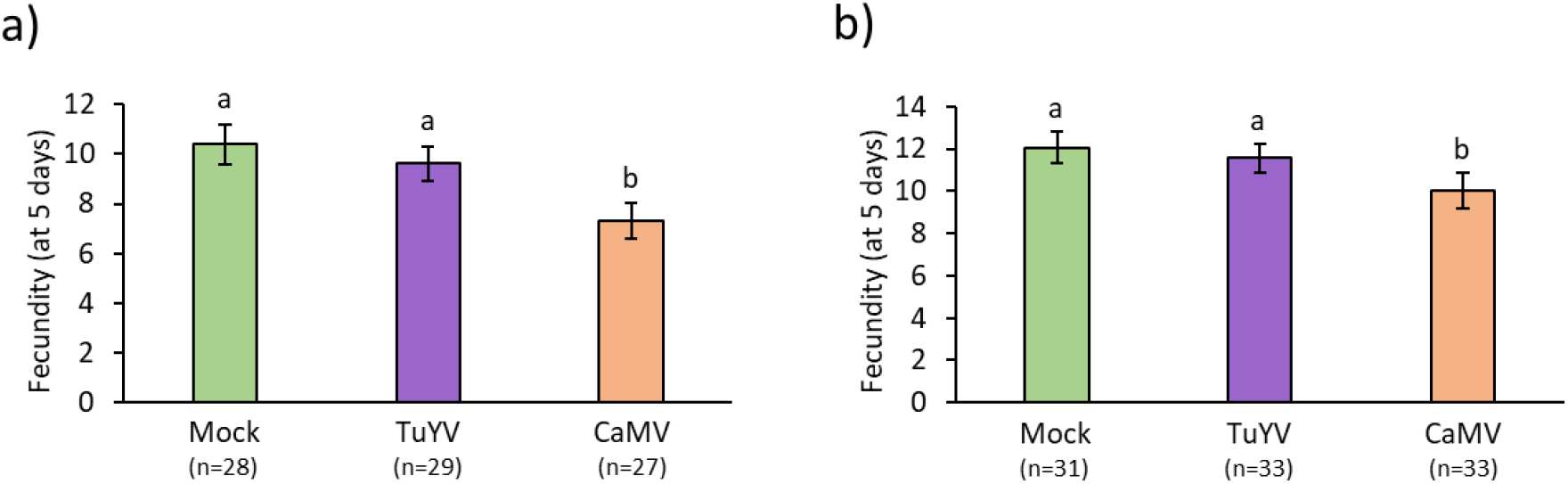
Aphid fecundity 5 days after deposit (one aphid per plant) on 5-week-old mock-inoculated, TuYV- or CaMV-infected a) Arabidopsis and b) Camelina. Different letters indicate significant differences between plants as tested by GLM followed by pairwise comparisons using “emmeans” (p < 0.05; method: Tukey, n = 27-33).

### Quality of RNA and sequence alignment data

Roughly 29-35 million reads were obtained for Arabidopsis mRNA-seq datasets, of which >80 % could be aligned for mock-inoculated and TuYV-infected samples and 80 % for CaMV-infected samples (Supplementary Table S2). For Camelina, 28-38million reads were obtained and 61-71 % of the reads could be aligned (Supplementary Table S2). Principal component analysis of RNA-seq libraries (Figure 4a,b) indicated for both plant species that the three biological replicates per condition clustered well together and that the different conditions (mock-inoculated or infected with either virus) were well separated. Thus, the reads were of excellent quality and suited for a transcriptome analysis.

**Figure 4:**
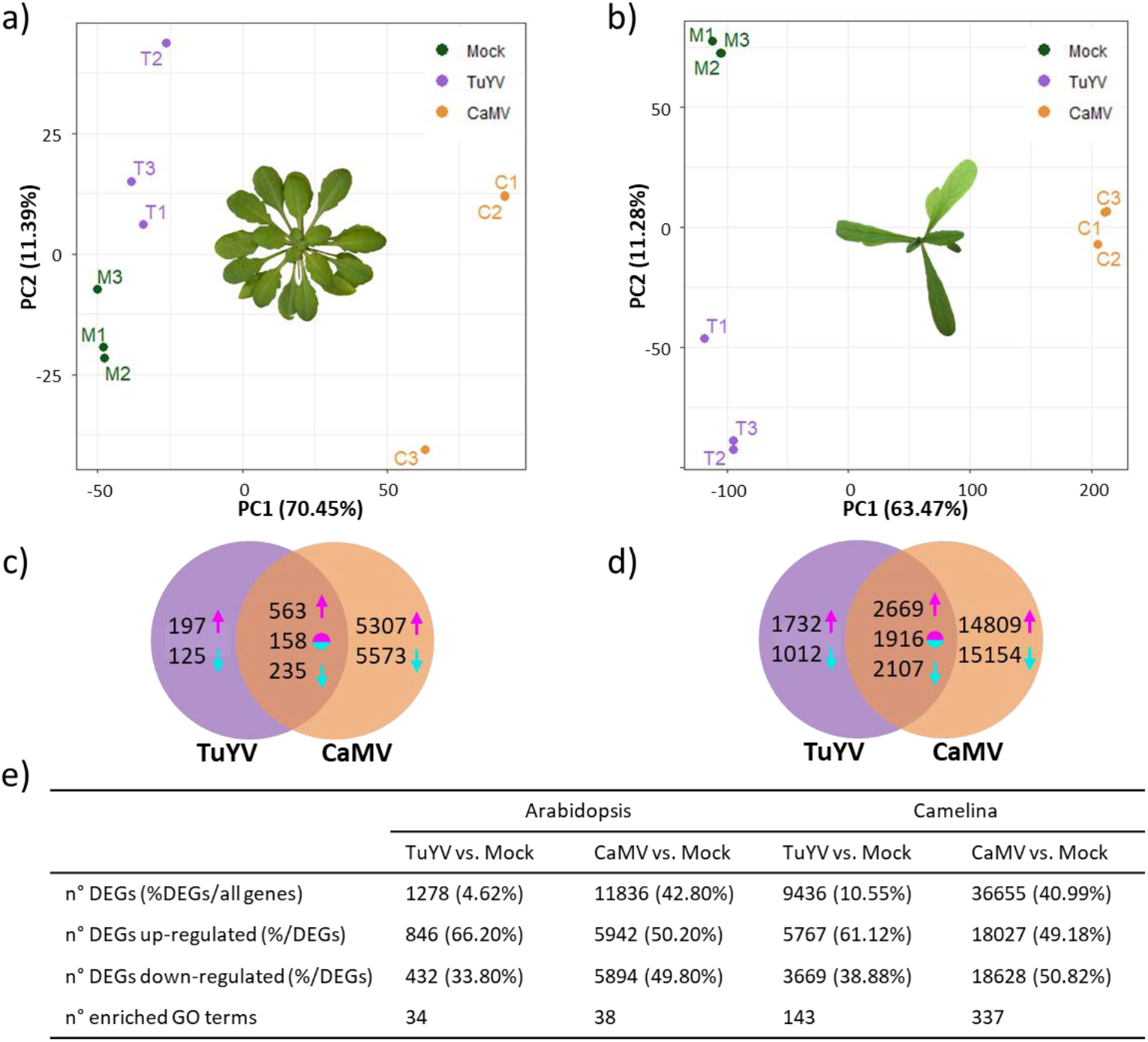
Principal component (PC) analysis of the transcriptome data sets on a) Arabidopsis and b) Camelina. The three dots of the same color correspond to the three biological replicates. (c and d) Venn diagrams presenting the number of differentially expressed genes (DEGs) in TuYV and CaMV-infected c) Arabidopsis and d) Camelina. Magenta arrows: number of up-regulated genes, cyan arrows: number of down-regulated genes and two-color circles: inversely regulated genes (up-regulated genes in one virus-infected modality and down-regulated in the other virus-infected modality). e) Comparison of the number of DEGs and enriched GO terms in TuYV and CaMV-infected Arabidopsis and Camelina plants.

For 8 selected Arabidopsis genes, the trends of gene deregulations detected in the transcriptome data could be reproduced by an alternative analysis method, RT-qPCR (Supplementary Figure S1 RT-qPCR). All 8 genes followed the same trend using either method for CaMV-infected Arabidopsis, and all except two (At_AOS and At_EDS5) for TuYV-infected Arabidopsis. The discrepancies were probably due to the rather weak expression changes, which are sometimes difficult to detect by RT-qPCR due to its intrinsic exponential amplification kinetics.

Quantification of viral RNA loads by counting viral reads normalized per million of total plant reads in each sample revealed that CaMV pregenomic (pg)RNA and TuYV genomic (g)RNA (both represented by forward reads; Supplementary Dataset S1) accumulated to comparable levels in each of the three biological replicates, with the exception of one of the three TuYV-Arabidopsis replicates which showed a lower number of normalized viral reads. The data confirmed further that the mock-inoculated plants were not cross-contaminated. Note that, because TuYV gRNA is not polyadenylated (unlike CaMV pgRNA) the poly(A) enrichment step of Illumina library preparation protocol should have led to its depletion. This might explain the greater variation in relative abundance of TuYV reads between biological replicates, compared to CaMV reads. Despite this high variability, a lower virus load was observed in TuYV-infected Arabidopsis compared to TuYV-infected Camelina plants (Supplementary Dataset S1). Notably, CaMV loads in Arabidopsis were also lower than those in Camelina (ca. 1.5 times; Supplementary Dataset S1). This indicates that despite drastic differences in disease symptoms between CaMV (severe symptoms) and TuYV (mild symptoms) in both Arabidopsis and Camelina, Camelina appears to be more conductive for replication of both viruses than Arabidopsis.

### CaMV modifies expression of far more genes than TuYV

Then we determined the number of differentially expressed genes (DEG) in infected hosts (Figure 4c-e). Far more DEGs were detected in Camelina than in Arabidopsis. This was in part due to its allohexaploid genome consisting of three Arabidopsis-like genomes coding for almost 90,000 genes (Kagale et al., 2014), compared to Arabidopsis’s diploid genome containing about 28,000 coding genes (http://ensembl.gramene.org/Arabidopsis_thaliana/Info/Annotation/#assembly, last accessed 17 December 2021). This means that for many Arabidopsis genes there are three orthologous Camelina genes. Also, the higher accumulation of both viruses in this host might contribute to the higher counts.

In Arabidopsis, CaMV modified expression of ~11,800 genes significantly (*P_(adj)_* <0.05), whereas TuYV modified expression of ~1,300 genes, corresponding to 43 % and 5 % of the total genes, respectively (Figure 4e). In CaMV-infected Camelina, we detected ~36,700 DEGs, and in TuYV-infected Camelina ~9,400 DEGs, corresponding to 41 % and 11 % of all genes, respectively. Thus, the impact of CaMV infection on gene deregulation was much more pronounced when compared to TuYV infection, in accordance with the phenotype of infected plants (Figure 1). The lower numbers of DEGs for TuYV in both hosts could be at least partially due to its restriction to phloem tissues, unlike CaMV which infects all cell types.

CaMV modified expression of ~40 % of the total genes independently of the host plant, whereas the proportion of TuYV-induced DEGs was host-dependent and two times higher in infected Camelina compared to Arabidopsis (11 % vs. 5 %). This is in line with the relative loads of viral RNA (Supplementary Dataset S1), indicating that Camelina is more susceptible to TuYV infection than Arabidopsis (3 times more TuYV RNA accumulation in Camelina compared to Arabidopsis), while CaMV accumulates in both hosts at comparable levels (only 1.5-fold difference in average viral RNA loads between Arabidopsis and Camelina).

956 DEGs, corresponding to 3.4 % of the genome, were common for both viruses in Arabidopsis. The proportion of common DEGs rose to 7.5 % (~6,700 genes) in infected Camelina. Since CaMV and TuYV are viruses with entirely different replication mechanisms, mediated by respectively viral reverse transcriptase and viral RNA-dependent RNA polymerase, these common host genes might be involved in general stress responses and/or are constituents of the core defense mechanisms. A GO analysis indicated that this was true for Arabidopsis with GO terms related to stress and transport in common for both infections, whereas for Camelina rather ribosome and replication-related genes were enriched (Supplementary Figure S2).

The proportions of up- and down-regulated genes were similar for a given virus in the two hosts (Figure 4e). However, when comparing the two viruses, the proportion of down-regulated genes was higher in CaMV-infected plants (about 50 % in CaMV-infected Arabidopsis and Camelina) than in TuYV-infected plants (34 % in TuYV-infected Arabidopsis 39 % in TuYV-infected Camelina). Thus, there was a correlation between the proportion of down-regulated genes and symptom severity. The milder disease symptoms of TuYV infection coincided in both hosts with the lower proportion of down-regulated genes, while the ability of CaMV to downregulate a higher proportion of genes coincided with the more severe disease symptoms. The latter ability might reflect CaMV activities in both cytoplasm (viral mRNA translation and pgRNA reverse transcription) and nucleus [viral dsDNA repair followed by pgRNA transcription and export assisted by nuclear-imported viral proteins P4, P5 and likely P6 (Haas et al., 2008; Kubina et al., 2021)].

### Impact of CaMV and TuYV infection on plant hosts: Gene ontology analysis

To identify the most prominent processes affected in aphid-infested CaMV and TuYV-infected Arabidopsis and Camelina, we carried out a Gene Ontology (GO) analysis (Figure 5). In general, TuYV-induced GO changes were much less pronounced (considering the percentage of DEGs in each category and the DEG counts) compared to CaMV, reflecting the low absolute numbers of DEGs in TuYV-infected plants and the weaker impact of TuYV on plant phenotype. Remarkably, in the Top 25 categories, only about 25 % of genes per GO were deregulated in TuYV-infected Arabidopsis (Figure 5a), whereas this value increased to more than 50 % in TuYV-infected Camelina (Figure 5c). Again, this may indicate that TuYV infection had a stronger effect on gene regulation in Camelina than in Arabidopsis. A different situation was found for CaMV, where the percentages of DEGs per GO were similar in both hosts, and always above 50 % of genes per GO, indicating similarly strong interactions with either host plant.

**Figure 5:**
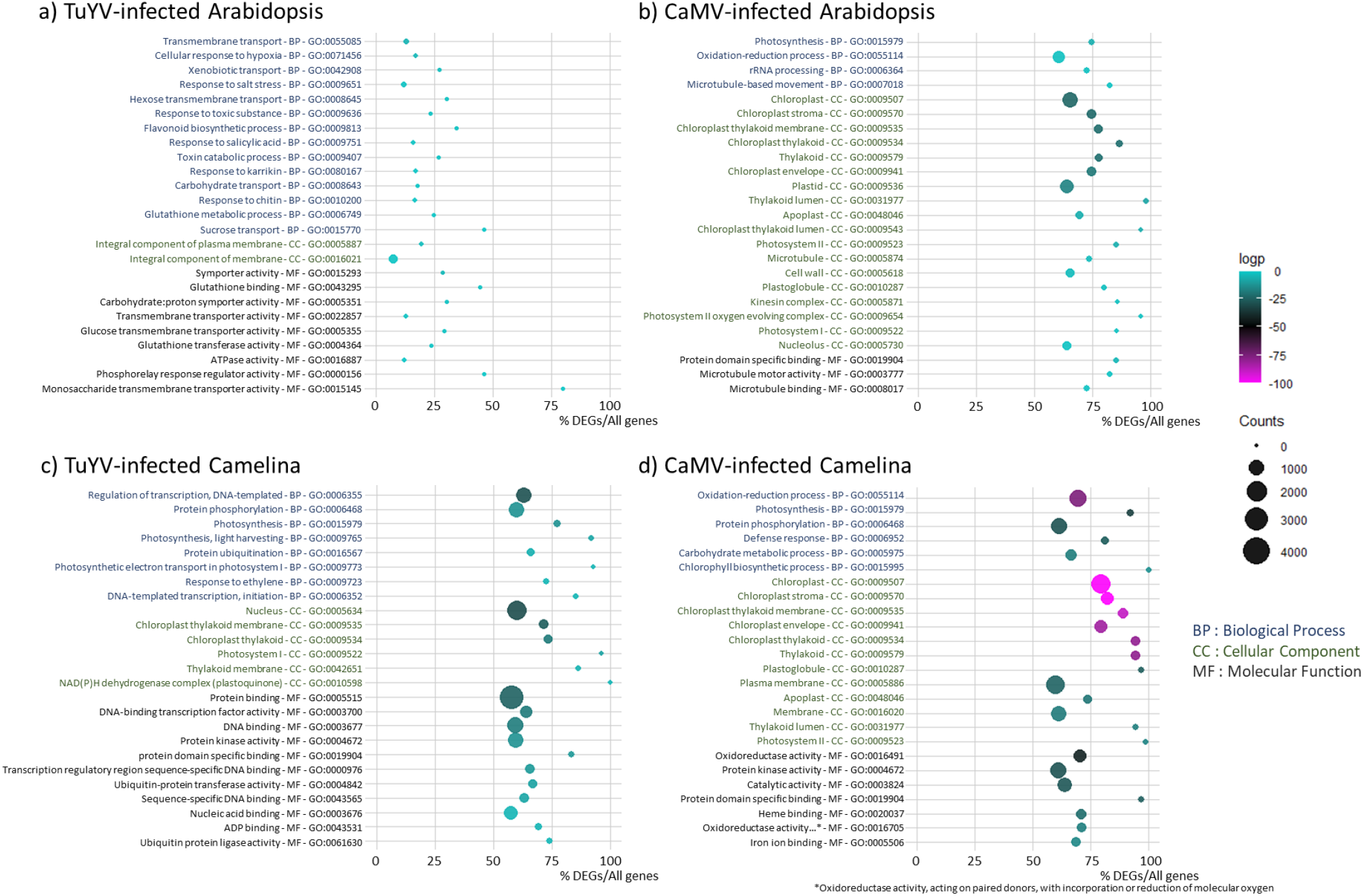
Gene ontology (GO) analysis showing the Top 25 GO of deregulated processes in TuYV- and CaMV-infected Arabidopsis and Camelina. a) TuYV-infected vs. mock-inoculated Arabidopsis, b) CaMV-infected vs. mock-inoculated Arabidopsis, c) TuYV-infected vs. mock-inoculated Camelina and d) CaMV-infected vs. mock-inoculated Camelina. GO IDs and corresponding GO terms are specified in the vertical axis. For each category (BP: Biological Process, CC: Cellular Component and MF: Molecular Function), GOs are sorted according to increasing log2 p-values, also indicated by the color of each spot (magenta representing the most significant p-values, see color scale bar), in order to place the most significantly enriched GOs on top of the graph. The absolute number of DEGs that matched the GO term is indicated by the size of each spot, whereas the horizontal axis shows the ratio of DEGs vs. all genes belonging to the GO term.

Then we looked closer at the different categories. Interestingly, in CaMV-infected Arabidopsis (Figure 5b), the majority of the enriched GO-terms was related to photosynthesis/chloroplast in both BP and CC categories, which might explain leaf chlorosis (loss of chlorophyll and/or chloroplasts). The next most affected biological process was oxidation-reduction that might be related to stress response. Also GO-terms related to microtubule-based movement appeared in the BP and CC lists, as well as apoplast, cell wall, kinesin complex and nucleolus which may be linked to virus or viral RNP intracellular trafficking. Taken together, CaMV infection mostly modified photosynthesis, oxidation-reduction processes and microtubule-related processes.

GO analysis of CaMV-infected Camelina indicated a similar pattern (Figure 5d). Again, several GO-terms related to photosynthesis/chloroplast and oxidation/reduction were enriched in both BP and CC categories. It is worth mentioning that for CaMV infection of Camelina, GO analysis showed enrichment of genes in the GO Defense Response (BP – GO:0006952), which was absent in the Top 25 list of CaMV-infected Arabidopsis. Other BP-related enriched GOs were protein phosphorylation and carbohydrate metabolism. As in Arabidopsis, apoplast changes were significant. On the other side, neither cell wall nor microtubule processes were present among the Top 25 deregulated processes in Camelina. Concerning the main categories of molecular functions, oxidation-reduction and protein domain specific binding dominated this category.

In contrast to CaMV, TuYV infection of Arabidopsis had no impact on photosynthesis-related GO terms. In both BP and MF, the most significantly enriched GO-terms were found in transport, especially carbohydrate transport. In addition, some defense and stress responses (xenobiotics, chitin, salt) and glutathione metabolism – indicative of oxidative stress – were affected. Flavonoid synthesis was significantly deregulated, in line with the purple-colored leaves of TuYV-infected Arabidopsis (Figure 1). In accordance with the modifications in sucrose transport, the most prominent category in CC comprised membranes. The Top 25 GO-terms in TuYV-infected Camelina were different from those in TuYV-infected Arabidopsis. As in CaMV infection, DEGs in photosynthesis and related processes dominated the Top 25 GO in TuYV-infected Camelina in BP and CC and were likely related to the mild yellowing symptoms appearing on old leaves. Next were DNA-related processes in both BP and MF categories, probably linked to transcriptional regulations of host genes in response to viral infection. In CC, the GO-term nucleus was deregulated, again in favor of a strong effect of TuYV on transcriptional regulation in infected Camelina. Other deregulated processes included ubiquitination, which appeared in several categories in BP and MF. On the other hand, oxidation-reduction did not appear under the top 25 GO except as plastoquinone, which represents a significant difference between CaMV and TuYV infections of Arabidopsis.

### Impact of CaMV and TuYV infection on plant hosts: Heatmap analyses

To better characterize the impact of viral infection on aphid-infested plants, we established the lists (Supplementary Dataset S2) and corresponding heatmaps (Figures 6-10) of DEGs for selected categories. Note that if not otherwise indicated, information on gene function is from the TAIR site (https://www.arabidopsis.org/). For mapping of Arabidopsis and Camelina genes in the heatmaps, we used the syntelog matrix (Kagale et al., 2014). This matrix assorts each individual Arabidopsis gene to the corresponding triplet of Camelina homologues. 62,277 Camelina genes out of 89,418 are syntenically orthologues (syntelogs) to Arabidopsis genes, among which some are considered ‘fractionated’ (if one or two of the homologues were lost). This explains why for certain Arabidopsis genes, only one or two homologues Camelina genes are presented.

**Figure 6.**
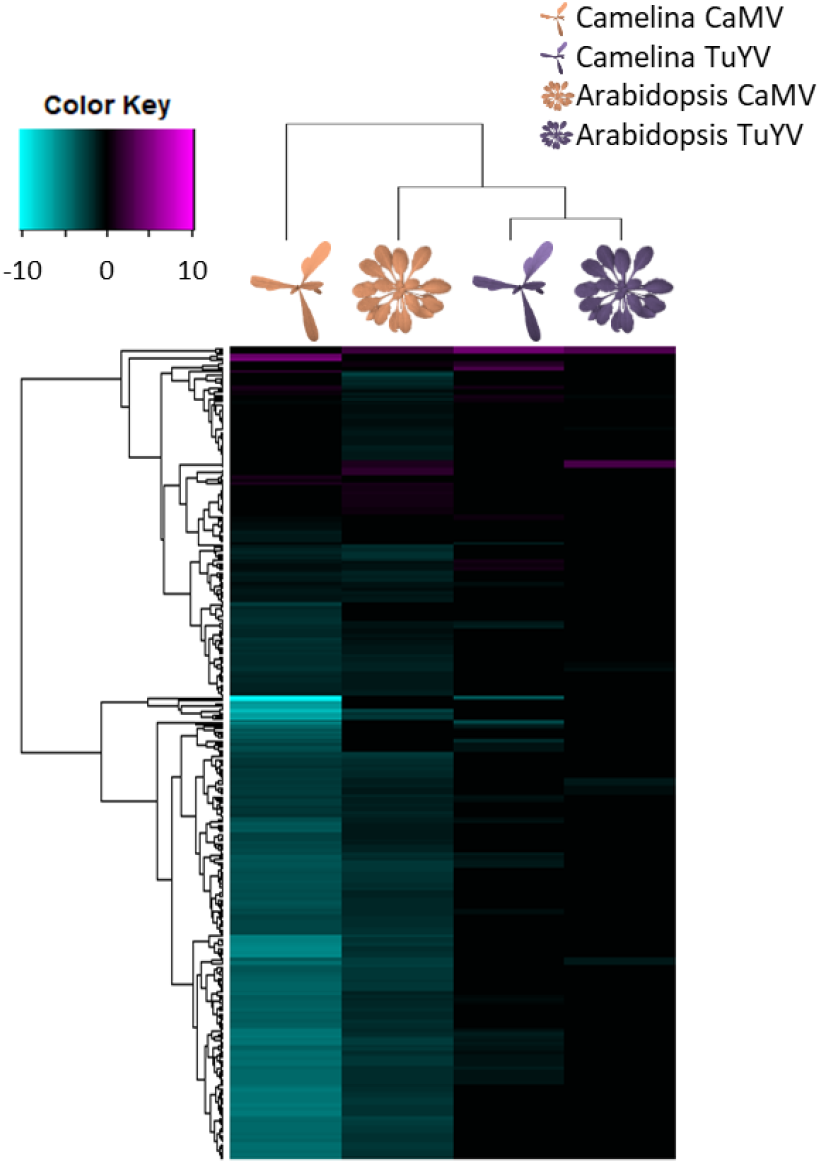
Hierarchical clustering of differentially expressed genes (DEGs) related to photosynthesis (GO:0015979) in CaMV- and TuYV-infected Arabidopsis and Camelina compared to their mock-inoculated control plants (Supplementary Dataset S2). The color key scale displays the log2fold changes from −10 to +10 as a gradient from cyan to magenta.

**Figure 7.**
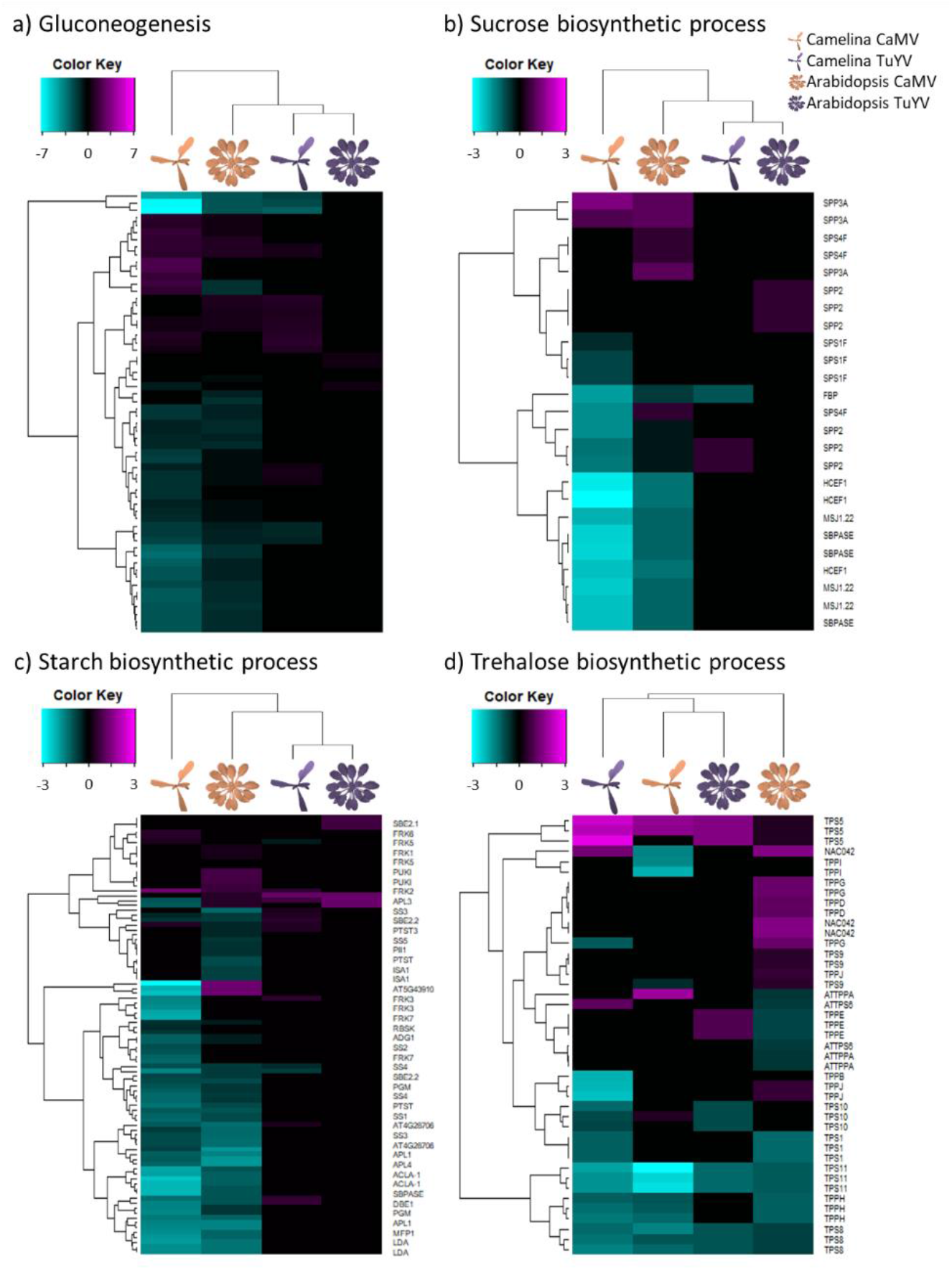
Hierarchical clustering of differentially expressed genes (DEGs) related to a) gluconeogenesis (GO:0006094), b) sucrose biosynthetic process (GO:0005986), c) starch biosynthetic process (GO:0019252) and d) trehalose biosynthetic process (GO:0005992) in CaMV- and TuYV-infected Arabidopsis and Camelina compared to mock-inoculated controls (Supplementary Dataset S2). The color key scales display the log2fold changes as color gradients from cyan to magenta.

**Figure 8.**
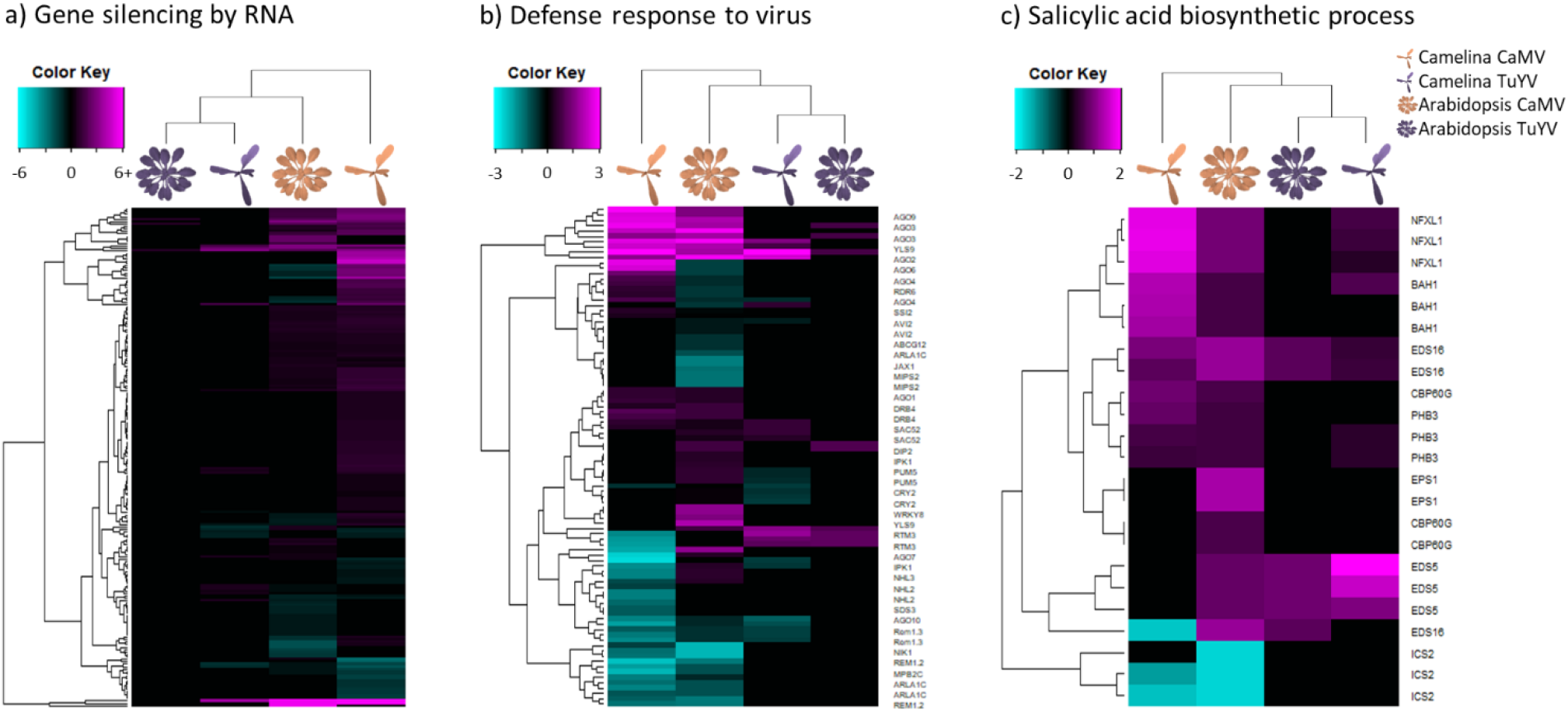
Hierarchical clustering of differentially expressed genes (DEGs) related to a) production of siRNA involved in RNA interference and Gene silencing by RNA (GO:0030422 and GO:0031047), b) defense response to virus (GO:0051607) and c) salicylic acid biosynthetic process (GO:0009697) in CaMV- and TuYV-infected Arabidopsis and Camelina compared to their mock-inoculated controls (Supplementary Dataset S2). The color key scales display the log2fold changes as gradients from cyan to magenta.

**Figure 9.**
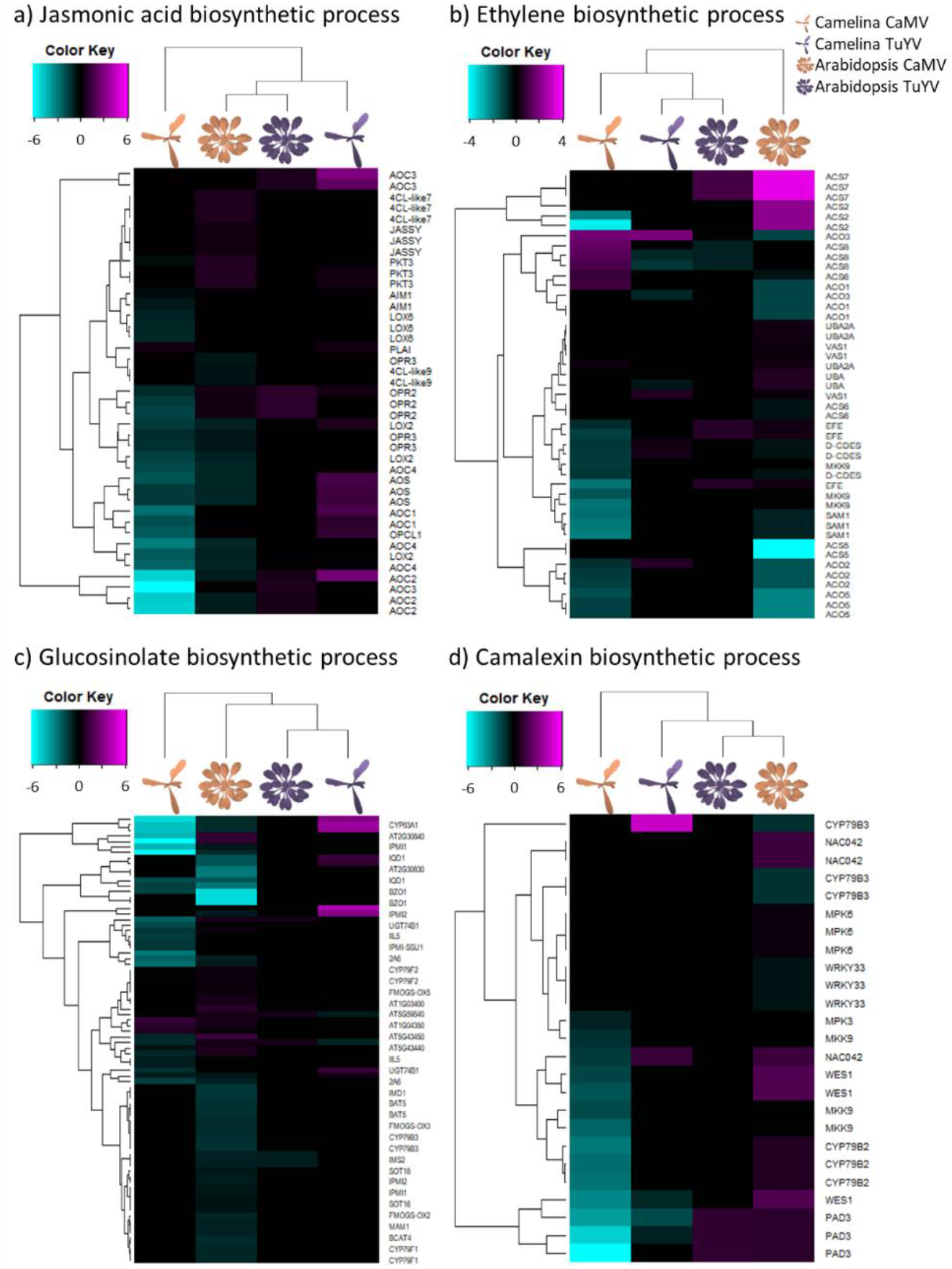
Hierarchical clustering of differentially expressed genes (DEGs) related to a) jasmonic acid biosynthetic process (GO:0009695), b) ethylene biosynthetic process (GO:0009693), c) glucosinolate biosynthetic process (GO:0019761) and d) camalexin biosynthetic process (GO:0010120) in CaMV- and TuYV-infected Arabidopsis and Camelina compared to mock-inoculated controls (Supplementary Dataset S2). The color key scales display the log2fold changes as gradients from cyan to magenta.

**Figure 10.**
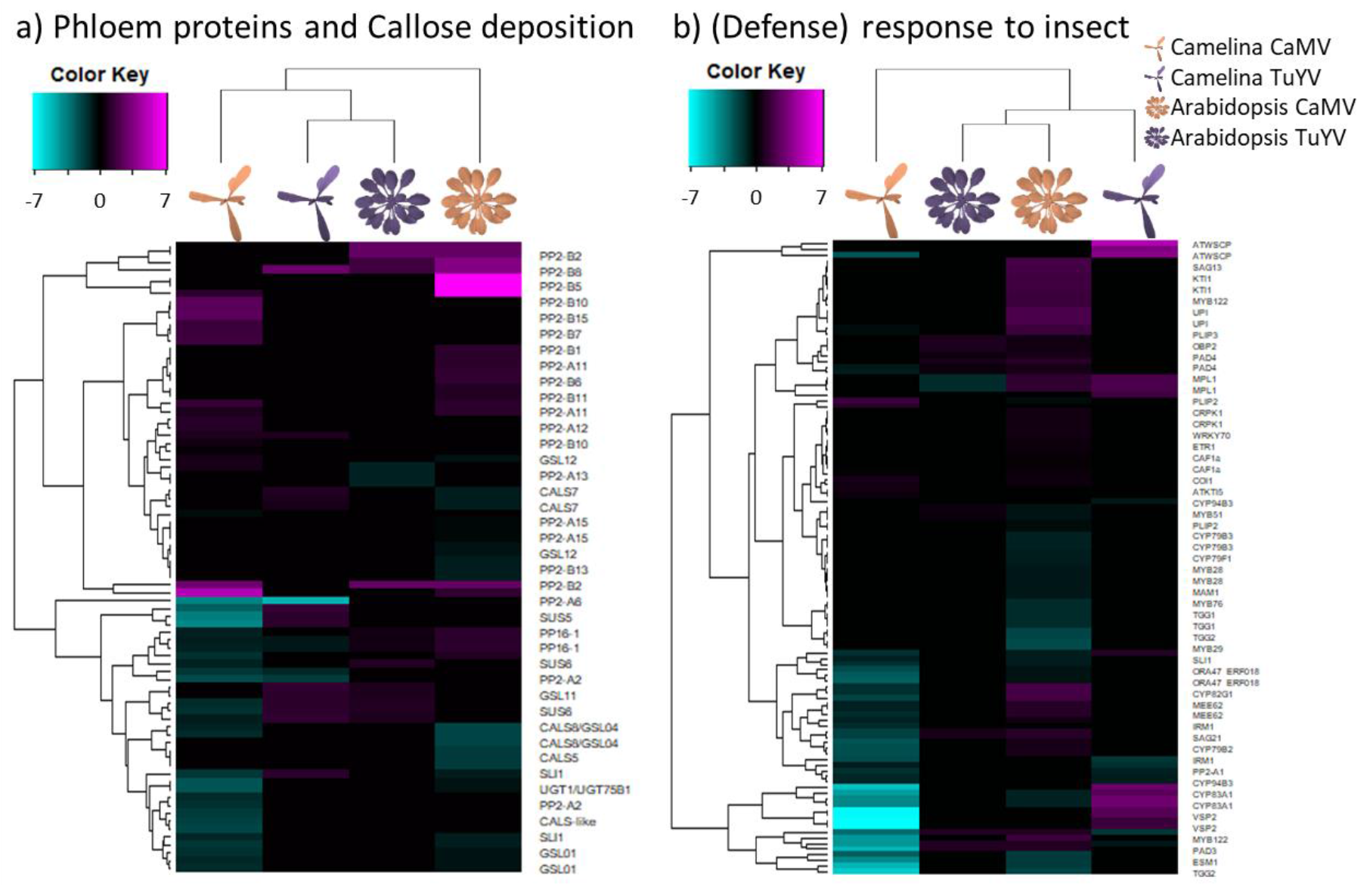
Hierarchical clustering of differentially expressed genes (DEGs) related to a) phloem proteins (PP2 and PP1) and callose deposition in phloem sieve plates (GO:0080165) and b) defense response to insect and response to insect (GO:0002213 and GO:0009625) in CaMV- and TuYV-infected Arabidopsis and Camelina compared to mock-inoculated controls (Supplementary Dataset S2). The color key scales display the log2fold changes as gradients from cyan to magenta.

CaMV and TuYV infection downregulated photosynthesis-related genes in infested Arabidopsis and Camelina (Figure 6). Overall downregulation of photosynthesis-related genes was much more pronounced in CaMV-infected than in TuYV-infected plants. Interestingly, both viruses interacted more strongly with Camelina photosynthesis than with Arabidopsis photosynthesis. No photosynthesis-related gene was similarly deregulated in all four conditions. The most downregulated photosynthesis gene in CaMV-infected Camelina was *PORA* (Camelina Csa02g051950, Csa11g086170 and Csa18g025480, corresponding to Arabidopsis AT5G54190), coding for a protein involved in chlorophyll biosynthesis, but also in response to ethylene. *PORA* expression was also inhibited in TuYV-infected Camelina. Expression of the Arabidopsis orthologue, however, was not modified by any of the two viruses. This might indicate that downregulation of *PORA* is a plant-specific and not a virus-specific response. The most downregulated gene in CaMV-infected Arabidopsis, AT3G27690, encoding the protein LHCb2.4, a component of the light harvesting complex, was also strongly repressed in Camelina infected by CaMV. Expression of this gene was also affected in TuYV-infected Arabidopsis but to a lesser extent and expression was not modified in TuYV-infected Camelina. Some genes were upregulated by infection. This was notably the case for the glucose-6-phosphate/phosphate transporter 2 (AT1G61800), which is involved in regulation of photosynthesis and which was upregulated in three of the four conditions (no gene expression modification in CaMV-infected Camelina). In CaMV-infected Camelina, a chloroplastic ferritin (AT3G11050) was the most upregulated photosynthesis gene. Ferritins are iron-binding proteins and are supposed to be involved in responses against oxidative stress in Arabidopsis (Briat et al., 2010), which could explain its overexpression.

Taken together, virus infection strongly interfered with photosynthesis. This might explain the leaf yellowing observed clearly on CaMV-infected plants and to a lesser extent on TuYV-infected Camelina. Leaf yellowing, probably due to reduced chlorophyll content in chloroplasts and/or to a reduced number of photosynthesizing chloroplasts caused by the gene deregulations (Chen et al., 2002), can alter settling preference of aphids (A’Brook, 1973; Fennell et al., 2020). However, this was not confirmed by previous observations on TuYV-infected and CaMV infected Camelina plants (Chesnais et al., 2019a). Indeed, although Chesnais and coworkers reported that *M. persicae* aphids preferred to settle on TuYV-infected Camelina, compared to healthy plants, no such aphid preference was observed for CaMV-infected Camelina, despite the strong yellowing symptoms. This suggests that aphid preference for a plant is not only driven by visual aspects.

In line with the repression of photosynthesis, expression of many sucrose synthesis and gluconeogenesis-related genes was reduced by CaMV infection (Figure 7a,b). The effect of CaMV was stronger in Camelina than in Arabidopsis. In TuYV-infected Camelina, the amplitude of the gene deregulation was smaller, compared to CaMV-infected Camelina, but the proportions of up- and down-regulated genes were comparable. In TuYV-infected Arabidopsis, expression changes were even smaller than in TuYV-infected Camelina. For both TuYV- and CaMV-infected plants, among the most down-regulated genes were those coding for key enzymes in sucrose synthesis, in particular *HCEF1* (AT3G54050) and *FBP* (AT1G43670). Interestingly, the sucrose phosphatase *SPP1* (AT1G51420) was strongly upregulated by CaMV infection, but downregulated in TuYV-infected plants. Remarkably, the most down-regulated gene in gluconeogenesis was the aldolase *FBA5* (AT4G26530) and this in all four conditions tested. In line with the stronger suppression of gluconeogenesis and sucrose synthesis-related genes by CaMV infection, many starch synthesis-related genes were also repressed by CaMV (but not TuYV) infection (Figure 7c). An exception was *DBE1* (Csa17g005380, Csa14g004380 and Csa03g004400, syntelogs of AT1G03310) encoding a starch branching enzyme was upregulated in TuYV-infected Camelina, which is consistent with a recent study showing that TuYV-infection tends to increase the carbohydrate concentrations in Camelina leaves (Chesnais et al., 2019b).

The effect of infection and infestation on trehalose metabolism was different from that on glucose, starch and sucrose metabolism (Figure 7d). Contrary to downregulation of the latter carbohydrate pathways only in CaMV-infected plants, trehalose-related genes were downregulated in both TuYV- and CaMV-infected Arabidopsis and Camelina. Upregulated genes in this pathway were also observed, in particular in CaMV-infected Arabidopsis. Trehalose is induced by *M. persicae* infestation and has been shown to contribute to defenses against aphids (Hodge et al., 2013; Singh et al., 2011; Singh and Shah, 2012). Especially *TPS11* (Trehalose Phosphate Synthase 11) has been implicated in mounting defenses against aphids (Singh et al., 2011), by promoting starch synthesis, but also by activating the phytoalexin-deficient gene, *PAD4.* Starch is a feeding-deterrent to aphids (Campbell et al., 1986) and elevated starch levels are correlated with reduced aphid performance (Singh et al., 2011). Interestingly, *TPS11* but also *TPS8* were significantly down-regulated in all virus-infected plants, suggesting that viral infection might favor aphid infestation. On the other hand, other TPS isoforms were not modified in the same way, for example the *TPS5* (AT4G17770) was upregulated in all four conditions tested. This suggests a more complex regulation of this pathway by viral infection and aphid infestation.

We also analyzed expression of genes involved in amino acid metabolism, because they represent the most important nutrient for aphids. Probably due to the multiple functions of these genes, no clear pattern was detected (Supplementary Figure S3).

When looking at global virus defense-related genes, the effects on their regulation were more virus-specific than host-specific. In agreement with previous findings in CaMV-infected Arabidopsis (Shivaprasad et al., 2008), many RNA silencing-related genes were found to be upregulated by CaMV not only in Arabidopsis but also in Camelina (Figure 8a). Among them, most notable are components of the 21 nt siRNA-directed gene silencing pathways such as Double-stranded RNA-binding protein 4 (DRB4), a partner of the antiviral Dicer-like protein 4 (DCL4) generating 21 nt siRNAs, and siRNA-binding effector proteins Argonaute 1 (AGO1; AT1G48410), AGO2 (AT1G31280) and AGO3 (AT1G31290). Notably, AGO2, also known to be involved in defense against RNA viruses (Carbonell and Carrington, 2015), was up-regulated in TuYV-infected Camelina (but not in TuYV-infected Arabidopsis), while AGO3 – the Argonaute protein most closely related to AGO2 and also showing antiviral activity *in vitro* (Schuck et al., 2013) – was up-regulated in response to TuYV infection in both hosts. This suggests redundant (and compensatory) roles of AGO2 and AGO3 in defenses against both RNA and DNA viruses. DCL4 itself (AT5G20320) and DCL2 (AT3G03300), generating 22 nt siRNAs and acting together with DCL4 in defenses against RNA and DNA viruses (Blevins et al., 2006), were respectively weakly (one of three isoforms of Camelina DCL4) and strongly upregulated (DCL2) in CaMV-infected Camelina but not in the other virus-host combinations where their levels were likely sufficiently high to cope with both viruses. Interestingly, AGO10 that counteracts AGO1 in the miRNA-directed silencing pathways regulating plant development and physiology (Yu et al., 2017) was downregulated by CaMV (but not TuYV) infection in both Arabidopsis and Camelina. RNA-directed RNA polymerase 6 (RDR6) generating miRNA-dependent secondary 21-nt siRNAs was upregulated by TuYV infection in Camelina and down-regulated by CaMV infection in Arabidopsis, while remaining non-responsive in the other virus-host combinations. Components of the nuclear silencing and 24 nt siRNA-directed DNA methylation pathways such as AGO4 (AT2G27040), AGO6 (AT2G32940) and AGO9 (AT5G21150) were upregulated in Camelina by CaMV (but not TuYV), while AGO4 and AGO6 were down-regulated and AGO9 up-regulated in CaMV-infected Arabidopsis, denoting virus- and plant-specific gene deregulations. Interestingly, the most upregulated gene in the RNA silencing category was AT5G59390. It was strongly upregulated in CaMV-infected Camelina and Arabidopsis, less strongly upregulated in TuYV-infected Camelina and not significantly deregulated in TuYV-infected Arabidopsis. This gene codes for a XH/XS domain-containing protein, which probably functions in siRNA-directed DNA methylation and might contribute to methylation and transcriptional silencing of CaMV dsDNA in the nucleus (Omae et al., 2020). Taken together, CaMV infection strongly affected silencing-related genes in both hosts, but the deregulations were host-specific, with down-regulations dominating in Arabidopsis, and upregulations in Camelina. Transcription of RNA silencing genes was much less affected in TuYV-infected plants.

Among components of other defense pathways (Figure 8b), the hairpin-induced protein hin1 (AT2G35980, also referred to as YLS9 and reported to be induced by cucumber mosaic virus infection) was strongly induced during CaMV infection, while Rem 1.2 (AT3G61260, also referred to as REMORIN and known to negatively regulate cell-to-cell movement of the potyvirus TuMV via competition with PCaP1 for binding actin filaments (Cheng et al., 2020) was strongly down-regulated during CaMV infection. None of these genes (*hin1* and *Rem1.2*) were deregulated by TuYV-infection. However, another REMORIN, Rem1.3 (known to impair potato virus X movement (Raffaele et al., 2009)) was down-regulated during TuYV infection in Camelina and during CaMV infection in both Arabidopsis and Camelina. On the other hand, the gene for myo-inositol-phosphate synthase 2 (*MIPS2*, AT2G22240) was downregulated in all conditions and *RTM3* [AT3G58350, known to block phloem movement of potyviruses, (Cosson et al., 2010)] was upregulated in TuYV- and downregulated in CaMV-infected hosts. It is therefore conceivable that remorins and MIPS2 are factors controlling TuYV and CaMV movement. Curiously, the gene *NIK1* (AT5G16000, NSP-interacting kinase), which encodes a receptor-like kinase, involved in innate immunity-based defense response against a ssDNA geminivirus, was strongly down-regulated in CaMV-infected but not in TuYV-infected host plants. Considering that downregulation of *NIK1* should activate protein translation and could promote accumulation of viral proteins (Zorzatto et al., 2015), it could have a proviral effect during CaMV infection.

Next, we looked at salicylic acid (SA) synthesis as this phytohormone is related to innate immunity-based defense responses against non-viral and viral pathogens including CaMV (Zvereva et al., 2016; Zvereva and Pooggin, 2012). Here, most genes were induced in both hosts and for both viruses (Figure 8c), with the notable exception of *ICS2* (AT1G18870), which was downregulated in CaMV infection and slightly upregulated in TuYV infection, while its redundant orthologue *ICS1* (AT1G74710) was slightly but significantly downregulated only in CaMV-infected Arabidopsis. Overall, genes involved in virus defense and SA biosynthesis were more strongly induced by CaMV-infection than TuYV-infection, whatever the host-plant, indicating a stronger pathogenicity of CaMV. The overexpression of SA-related genes in CaMV-infected plants could also reflect the ability of CaMV effector protein P6 to suppress SA-dependent autophagy, which might lead to compensatory feedback up-regulation of SA genes (Zvereva et al., 2016). Deregulation of genes implicated in the SA pathway might have consequences on insect-plant interactions. In particular increased SA could have a beneficial effect on aphid vector and possibly transmission, because it can be concomitant with decreased JA levels and consequently with decreased plant defenses against aphids (Kloth et al., 2016; Lu et al., 2020).

Next, we analyzed different metabolic pathways to determine if CaMV and TuYV infections modulate other hormones and secondary metabolites in ways that are more favorable for their aphid vector, and hence, for their own transmission.

We first looked at jasmonic acid (JA) and its derivatives because they are plant signaling molecules related to plant defense against herbivorous insects, microbial pathogens and different abiotic stresses (Figure 9a). We observed a strong virus-specific and host-independent effect for JA synthesis genes. CaMV downregulated many genes in the JA pathway, while TuYV upregulated some. Like for other pathways, the effect was stronger in infected Camelina than in Arabidopsis. Deregulated genes were for example *AOC1/3/4* (3 out of for 4 chloroplastic allene oxide cyclases, involved in JA synthesis), *AOS* (AT5G42650, chloroplastic allene oxide synthase, involved in JA synthesis) and *LOX2* (AT3G45140, chloroplastic lipoxygenase required for wound-induced JA accumulation in Arabidopsis). All these genes were slightly upregulated in TuYV, and strongly repressed in CaMV-infected plants. This might imply that JA production is down in CaMV-infected plants and stable or slightly induced in TuYV-infected plants. JA is generally thought to decrease aphid growth and fecundity, so aphids on CaMV-infected plants might have greater fecundity. However, infection of Arabidopsis with CaMV lowered fecundity (Figure 3a). JA-mediated signaling pathways are also known to increase proteins and secondary metabolites, which act as feeding deterrents (Howe and Jander, 2008). In this context, decreased JA production in CaMV-infected Arabidopsis could encourage longer/faster phloem sap ingestion, which we observed indeed in our experiments (Figure 2a). Interestingly, phloem sap ingestion has been correlated with increased CaMV acquisition (Palacios et al., 2002), which makes JA pathway a major candidate for virus manipulation.

We also analyzed ethylene (ET) synthesis (Figure 9b) as several studies have identified ET as a plant response against aphid infestation (Anstead et al., 2010; Mewis et al., 2005). No specific gene expression patterns characteristic for a virus or a host were found, indicating that the ethylene response was unique for each virus-host pair. Noteworthy, the ACC oxidase genes *ACO2* and *ACO5* (AT1G62380 and AT1G77330), involved in ethylene production, were strongly down-regulated for CaMV-infected host-plants. This is consistent with reduced accumulation of ethylene in CaMV-infected and P6-transgenic Arabidopsis in response to bacterial elicitors of innate immunity (Zvereva et al., 2016).

Glucosinolates (GLSs) are secondary metabolites that are produced by plants in the *Brassicaceae* family and set free in response to herbivore attacks (Kim et al., 2008). Some GLSs have been shown to be strong feeding deterrents for generalist aphids such as *M. persicae* (Kim and Jander, 2007) and might even have antibiosis effects on this aphid species (Cole, 1997; Westwood et al., 2013). CaMV infection down-regulated genes involved in GLS synthesis (Figure 9c), for example, the three Camelina orthologues encoding the cytochrome P450 monooxygenase CYP83A1 (AT4G13770), whereas these three genes were upregulated in TuYV-infected Camelina. The effect of CaMV infection on CYP83A1 was less pronounced in Arabidopsis, where another gene, AT1G65880, involved in benzoyloxyglucosinolate 1 synthesis, was strongly repressed. Other genes implicated in GLS synthesis, for example *IMD1* (AT5G14200), an isopropylmalate dehydrogenase (He et al., 2011) were upregulated in TuYV-infected Camelina. The transcription factor *MYB51* (AT1G18570), involved in indole glucosinolate synthesis (Barco and Clay, 2020), was downregulated in CaMV-infected Camelina, but not in the other conditions. All in all, infection with CaMV predominantly down-regulated transcription of GLS-related genes in Camelina and to a lesser extent in Arabidopsis, whereas TuYV infection induced GLS synthesis in aphid-infested Camelina, and had hardly any effect on Arabidopsis. We therefore expected that *M. persicae* fitness and feeding behavior (i.e. phloem sap ingestion and ease to access phloem tissues) would be enhanced on CaMV-infected Camelina and Arabidopsis, and be decreased on TuYV-infected Camelina. However, our fecundity experiments show that, on the contrary, *M. persicae* fecundity was decreased on CaMV-infected Arabidopsis and remained unchanged on TuYV-infected Arabidopsis compared to mock-inoculated plants, while no effects were observed on Camelina plants. Previous experiments indicated that *M. persicae* fecundity is even higher in TuYV-infected Camelina and lower in CaMV-infected Camelina (Chesnais et al., 2019a). Note, however, that in the experiment of Chesnais et al. (2019a), a more severe CaMV strain was used, which might explain the discrepancies between both experiments. Overall, based on our results, deregulation of GLS-related genes after CaMV and TuYV infections do not seem to be the main factors controlling *M. persicae* fecundity. This is in line with another study that found no correlation between the GLS content of rapeseed and *M. persicae* fecundity (Weber et al., 1986).

On the other hand, our EPG experiments showed that aphids were able to reach phloem tissues and ingest phloem sap for a longer duration on CaMV-infected Arabidopsis and Camelina (see also Chesnais et al., 2019a; Chesnais et al., 2021). Therefore, down-regulation of GLS genes might encourage aphid settling/feeding behavior on CaMV-infected plants, and eventually promote CaMV acquisition by *M. persicae*. On TuYV-infected plants, while some GLS-related genes were slightly up-regulated, aphid feeding behavior was roughly equivalent to that on healthy plants, indicating that up-regulations were not strong enough to induce feeding deterrence.

Camalexin is the major phytoalexin and has been shown to reduce fecundity of aphids in Arabidopsis (Kettles et al., 2013), although its effect on aphids might not be straight-forward (Kloth et al., 2019; Pegadaraju et al., 2005). PAD3 (phytoalexin deficient 3) catalyzes the last step in its synthesis and CYP79B2 an important intermediate step. Here we found contrasting effects of aphid infestation on virus-infected plants on camalexin-related genes (Figure 9d). *PAD3* was downregulated in CaMV-infected and to a lesser extent in TuYV-infected Camelina, and slightly upregulated in Arabidopsis infected with CaMV or TuYV. *CYP79B2* expression was unaffected in both Camelina and Arabidopsis infected with TuYV, but substantially downregulated in CaMV-infected Camelina and slightly upregulated in CaMV-infected Arabidopsis. This indicates for both genes a strong host-plant effect. Evidence indicates that PAD3 contributes more to camalexin synthesis than CYP79B2 (Kim et al., 2015; Zang et al., 2008; Zhang et al., 2020). Thus, looking at *PAD3*, aphid fecundity should be higher on CaMV- and TuYV-infected Camelina, and lower on infected Arabidopsis. However, we observed a lower fecundity in CaMV-infected Arabidopsis and unchanged aphid fecundity in all other conditions, which suggests that aphid fecundity is not only linked to *PAD3* expression. On the other hand, phloem ingestion on both CaMV-infected Arabidopsis and Camelina increased which is more in accordance with aphid plant acceptance. Overall, camalexin-related gene deregulations observed in both infected host plants did not seem to correlate with modified aphid fecundity, nor with aphid feeding behavior.

Callose is a polymer that is deposited by plants in between cells and in sieve tubes to restrict access of pathogens, including aphids, to tissues and phloem (Kuśnierczyk et al., 2008). We did not find any major DEG for this category except the stress-related plasma membrane respiratory burst oxidase Rboh F (Suzuki et al., 2011) and the pectin methylesterase inhibitor AT5G64640. No clear pattern of gene deregulation was observed, making interpretation difficult (Supplementary Figure S2). This might also be due posttranslational modifications that majorly regulate RbOH F activity (Kadota et al., 2015).

Since aphid lifestyle depends on compatible interactions with the phloem they feed on, we looked at phloem protein expression (Figure 10a). In all conditions except aphid-infested TuYV-infected Camelina, *PP2-B2* was the strongest induced gene. *PP2-B2* codes for a phloem-specific lectin-like protein with unknown function containing an F-box domain and a potential myristoylation site (Boisson et al., 2003) that could control membrane localization. Specific for CaMV infection, the putative phloem lectin genes *PP2-B1* (AT2G02230) and *PP2-B5* (AT2G02300) were upregulated in Arabidopsis and one of their orthologues was upregulated in Camelina. The putative calcium-, lipid- and RNA-binding phloem protein PP16-1 (AT3G55470) was, independent of the virus, upregulated in infected Arabidopsis and downregulated in Camelina. One Camelina orthologue of the Arabidopsis PP2-A1, known to repress aphid phloem feeding (Zhang et al., 2011), was down-regulated in Camelina infected with both CaMV and TuYV, but in Arabidopsis this gene remained non-responsive to viral infections. It is worth mentioning that PP2 proteins of cucurbits could potentially bind to viral particles of CABYV (genus *Polerovirus* like TuYV) and increase virus stability in the aphid gut (Bencharki et al., 2010). Proteins of this type could therefore have a double importance due to their role on vector aphids’ feeding behavior and their possible involvement in virus transmission. Deregulation of most other phloem proteins did not follow a distinct pattern and the unknown functions of most of these genes precluded any interpretation.

CalS7 (AT1G06490), a phloem-specific callose synthase responsible for wounding stress-induced callose deposition onto sieve tube plates and hence phloem plugging (Xie et al., 2011), was slightly upregulated in TuYV-infected Camelina and downregulated in CaMV-infected Arabidopsis. The same trend (upregulation in TuYV-infected Camelina and Arabidopsis, downregulation in CaMV-infected Camelina) applied to the phloem-located sucrose synthases SUS5 and SUS6 (AT5G37180 and AT1G73370) that interact with CalS7 (Barratt et al., 2011). Also *SLI1* (AT3G10680), a gene coding for a phloem small heat shock-like protein known to be involved in resistance to *M. persicae* and other phloem feeders (Kloth et al., 2021), was downregulated in both Arabidopsis and Camelina infected by CaMV. This might indicate that CaMV infection but not TuYV infection favors phloem feeding of aphids by perturbing stress-related callose deposition on sieve plates. This is in line with the prolonged phloem ingestion observed for *M. persicae* on CaMV-infected plants (Figure 2).

Next, we examined expression of genes known to be involved in plant responses and defenses against insects (Figure 10b), as their modulation could influence virus-insect interactions and hence transmission. General trends were suppression in CaMV-infected Camelina and activation in CaMV-infected Arabidopsis and in TuYV-infected Camelina and Arabidopsis, resulting in both host-specific and virus-specific responses. *ESM1* (AT3G14210) was strongly downregulated in both CaMV-infected hosts, but not in TuYV-infected hosts. Its gene product biases production of glucosinolates, and its knockout mutant is more susceptible to herbivory by the caterpillar *Trichoplusia ni* (Zhang et al., 2006). Thus, its downregulation in CaMV-infected plants might favor aphid colonization. Expression of ATWSCP (AT1G72290), a protease inhibitor and water-soluble chlorophyll-binding protein, was strongly upregulated in TuYV-infected and downregulated in CaMV-infected Camelina, whereas its expression was unchanged in Arabidopsis. The apoplastic ATWSCP, together with the protease RD21, protects plants, especially greening plants, against herbivory (Boex-Fontvieille et al., 2015). Whether it also acts against aphids is unknown. The *M. persicae*-induced lipase 1 (MPL1, AT5G14180) was upregulated in TuYV-infected Camelina and CaMV-infected Arabidopsis, but downregulated in TuYV-infected Arabidopsis and unaffected in CaMV-infected Camelina. This gene is induced by aphid infestation and decreases aphid fecundity, but it does not change aphid behavior or plant choice (Louis et al., 2010). Whether the reduced fecundity of *M. persicae* on CaMV-infected Arabidopsis is partially due to the action of this gene, remains an open question. Strong host plant-specific and virus-specific effects were found for VSP2 [AT5G24770, reported to have a role in defense against herbivory insects (Liu et al., 2005)], whose expression was up-regulated in aphid-infested TuYV-infected Camelina and down-regulated in aphid-infested CaMV-infected Camelina but not affected in infested Arabidopsis. All in all, plant defense responses against insects did not follow a clear pattern. This was probably due to the very divergent pathways and the heterogeneity of the plant insect response genes.

### Concluding remarks

In this work we analyzed the effect of CaMV and TuYV infection of *M. persicae* aphid-infested Arabidopsis and Camelina on the plant hosts’ transcriptomes as well as on the fecundity and feeding behavior of their vector *M. persicae*.

Our results show that CaMV infection caused more severe effects on phenotype of both plant species than did TuYV infection (Figure 1). The severity of symptoms correlated strongly with the proportion of DEGs (41-43 % for CaMV, 5-11 % for TuYV, Figure 4e). CaMV infection affected the same percentage of genes in both plant hosts, whereas TuYV infection deregulated proportionally twice as much genes in Camelina than in Arabidopsis. Again, this correlated with stronger visible symptoms on TuYV-infected Camelina in comparison with TuYV-infected Arabidopsis. Aphid performance changes were more pronounced on CaMV-infected hosts, whatever the plant species, compared to those caused by TuYV infection. In spite of more DEGs in TuYV-infected Camelina than in TuYV-infected Arabidopsis, aphid behavior was slightly more impacted on TuYV-infected Arabidopsis (Figure 2). This likely indicates modification of plant metabolites that cannot be identified by transcriptome profiling. A metabolomic analysis of virus-infected leaves or phloem sap should provide complementary data on the aphid-plant-virus interactions.

In this study, we did not compare the contribution of aphid infestation alone on the plant transcriptome. However, recent work (Annacondia et al., 2021) on the transcriptome changes of healthy Arabidopsis plants infested or not with *M. persicae* for 72 h, identified a limited number of DEGs (265) suggesting that the contribution of aphid infestation to the transcriptome in healthy and probably also in virus-infected and infested plants is minor.

The most pronounced effect of CaMV infection on plant hosts was a strong downregulation of photosynthesis genes (Figure 6) and carbohydrate metabolism-related genes (Figure 7). We observed significant changes in many other pathways, including categories that are likely affecting virus-vector interactions (*i.e.* defenses, silencing, hormones, secondary metabolites etc.). However, the impact of these modifications on aphid fitness or feeding behavior was not easy to evaluate since these parameters are likely under the control of several, often overlapping metabolic pathways. Trying to correlate the effect of specific genes on aphids as reported in the literature with our aphid behavior observations therefore often resulted in contrasting results. We offer the following explanations. The very strong alterations in photosynthesis might have drowned otherwise visible effects of DEGs previously found to be involved in plant-aphid interactions. Another explanation are posttranscriptional and posttranslational modifications. While transcriptome profiling is a powerful tool, it can display only changes of transcript levels. In many cases, however, posttranslational modifications of proteins (such as phosphorylation, localization, complex formation and many more) and even posttranscriptional RNA modifications (sequestering of RNAs in p-bodies and others) will contribute to phenotype changes. Depending on the pathway, the contribution of transcriptome and posttranscriptome on cellular processes and beyond will vary. This again indicates that complementary analyses such as metabolomics, proteomics etc. might help to gain a more complete insight.

Nevertheless, we observe that virus infections, whatever the host plant, have very distinct effects on the transcriptome of host plants, and that, as expected, the non-phloem-limited virus (i.e. CaMV) has a significantly stronger impact on plant hosts than the phloem-limited virus (i.e. TuYV). Overall, viral infection with CaMV tends to have effects on metabolic pathways with strong potential implications for insect-vector / plant-host interactions, while TuYV only weakly alters these pathways. For example, the strong gene downregulations in the jasmonic acid, ethylene and glucosinolate biosynthetic processes (Figure 9a-c) in CaMV-infected plants could be responsible for the observed alterations of aphid feeding behavior and performances. Next steps could consist in functional validation of some candidate genes identified in our study for their role in viral manipulation and consequently potential impacts on viral transmission.

## Supporting information

Supplementary Data Sets and Sequence Information

## Data availability

The raw RNA-seq data are available under project number PRJEB49403 at the European Nucleotide Archive (https://www.ebi.ac.uk/ena/browser/view/PRJEB49403).

## Author contributions

Conceptualization, Q.C., V.B., M.P. and M.D.; methodology, Q.C., V.G. and M.D.; software, Q.C., A.V. and C.R.; validation, Q.C. and V.G.; formal analysis, Q.C., A.V., C.R. and M.D.; investigation, Q.C. and V.G.; Data curation, Q.C., A.V. and C.R.; Writing – Original Draft Preparation, Q.C., M.P. and M.D.; Writing – Review & Editing, Q.C., A.V., C.R., V.B., M.P. and M.D.; Visualization, Q.C.; supervision, M.P. and M.D.; project administration, M.D.; funding acquisition, M.P. and M.D.

## Acknowledgments

We thank Claire Villeroy for aphid rearing and the experimental unit of INRAE Grand Est – Colmar (UEAV) for help with plant production. This work was supported by a public grant overseen by the French National Research Agency (ANR) (reference: ROME ANR-18-CE20-0017-01). Dr. Quentin Chesnais was supported by Région Grand Est (Soutien aux jeunes chercheurs, reference: 18_GE5_013). The funding sources had no role in the study design; in the collection, analysis, and interpretation of data; in the writing of the report; and in the decision to submit the article for publication. The authors declare no conflict of interest.

## Supplemental data

Provided as extra files:

Supplementary Dataset S1 Plant mRNA-seq.xlsx

Supplementary Dataset S2 Heatmaps DEGs List.xlsx

Supplementary Sequence Information S1 on CaMV and TuYV.docx

**Table S1:**
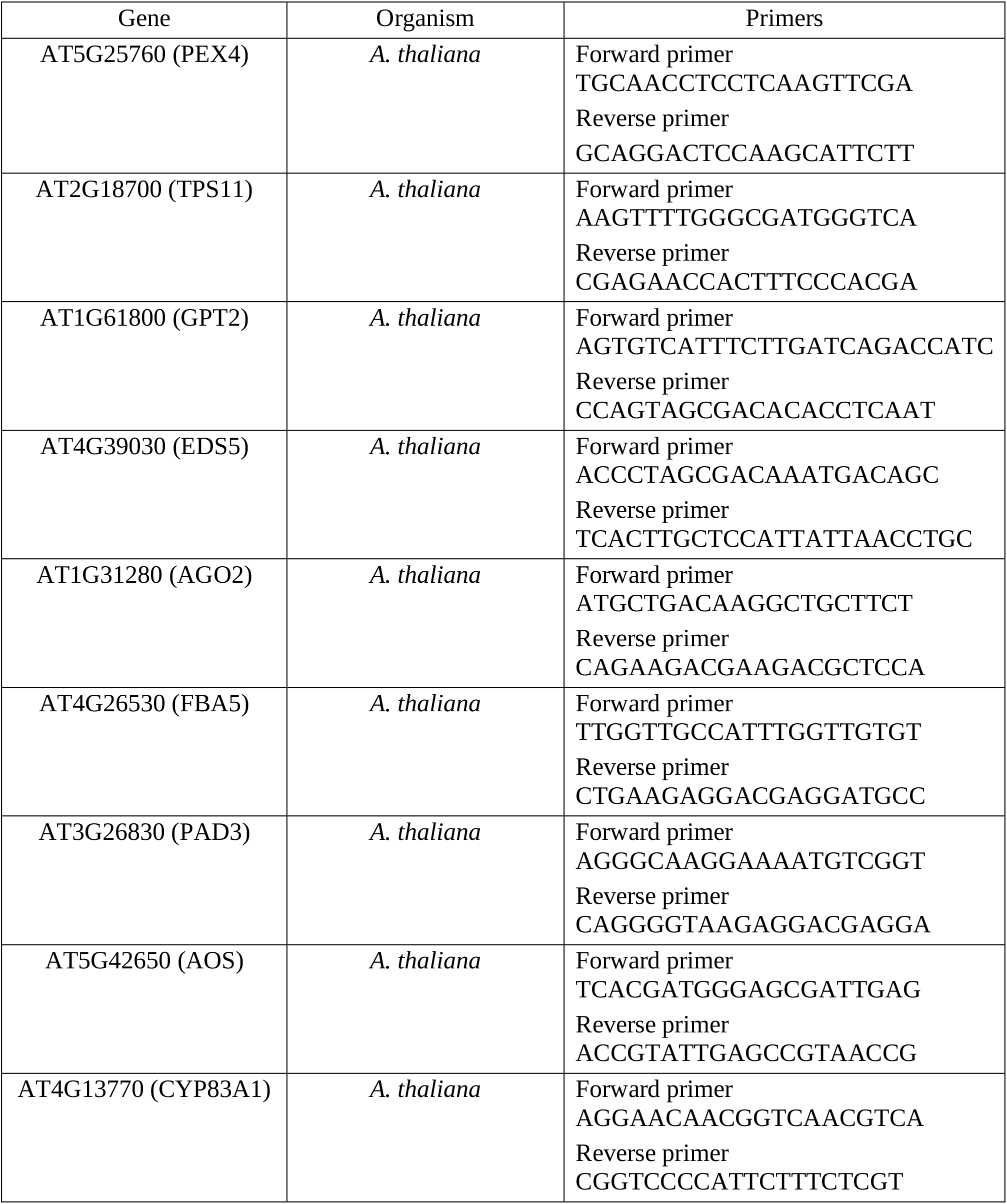
Oligonucleotides used for RT-qPCR

**Table S2:** Aligned reads for transcriptome profiling

| Sample name | Aligned reads | Assigned Reads | Mapped Ratio |
| --- | --- | --- | --- |
| Ara_M2 | 34,670,703 | 31,998,776 | 92,3% |
| Ara_M3 | 31,050,325 | 28,078,404 | 90,4% |
| Ara_M4 | 29,883,629 | 26,930,049 | 90,1% |
| Ara_C1 | 33,833,716 | 27,677,748 | 81,8% |
| Ara_C2 | 30,911,854 | 24,741,132 | 80,0% |
| Ara_C3 | 30,653,084 | 24,528,34 | 80,0% |
| Ara_T1 | 29,290,278 | 25,109,740 | 85,7% |
| Ara_T2 | 32,381,425 | 28,898,406 | 89,2% |
| Ara_T3 | 32,197,738 | 28,344,993 | 88,0% |
| Cam_M1 | 33,109,309 | 22,574,696 | 68,2% |
| Cam_M2 | 32,233,737 | 22,145,697 | 68,7% |
| Cam_M3 | 33,890,114 | 22,984,537 | 67,8% |
| Cam_C1 | 38,817,320 | 24,936,094 | 64,2% |
| Cam_C2 | 31,267,901 | 20,165,623 | 64,5% |
| Cam_C3 | 28,166,913 | 17,126,400 | 60,8% |
| Cam_T1 | 29,673,896 | 20,168,532 | 68,0% |
| Cam_T2 | 29,764,440 | 20,762,349 | 69,8% |
| Cam_T3 | 34,740,771 | 24,509,728 | 70,6% |

**Figure S1.**
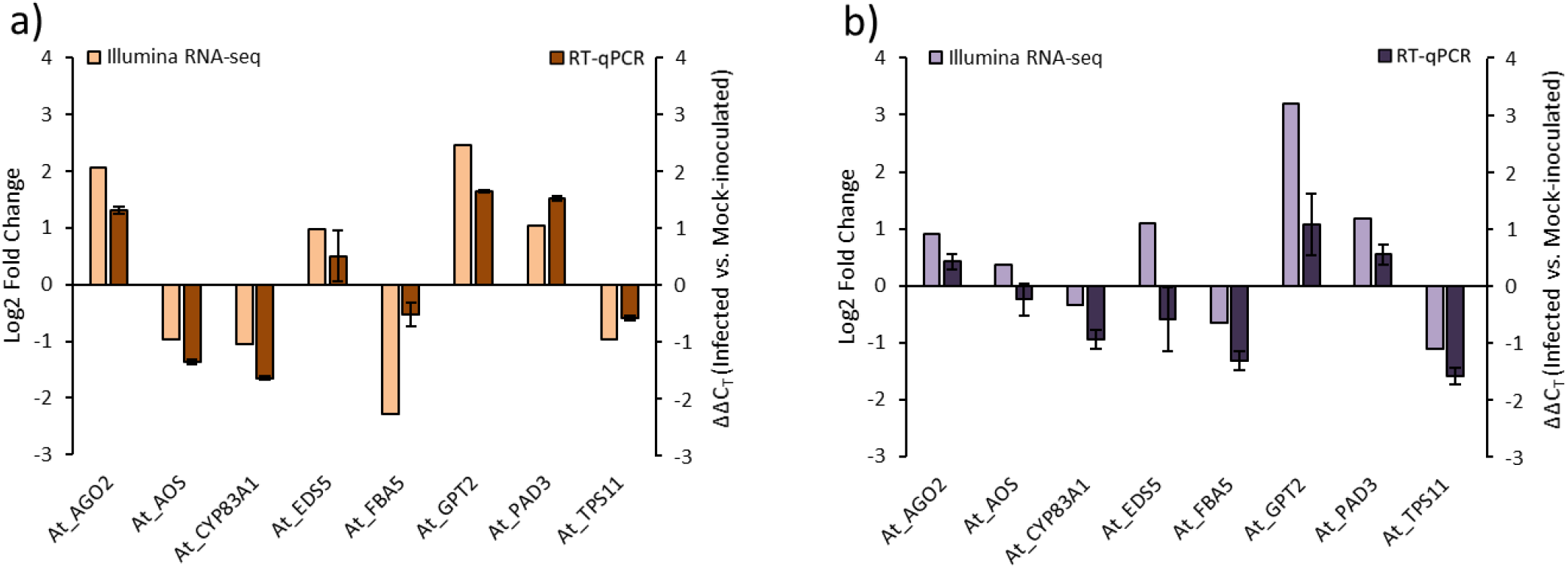
Validation of Illumina RNA-seq expression data by quantitative reverse-transcription PCR (RT-qPCR). a) CaMV-infected Arabidopsis. b) TuYV-infected Arabidopsis. The y-axis presents the normalized log2 fold change of expression derived from Illumina RNA-seq read counts and PCR ΔΔC_T_, respectively. The TAIR gene loci of the tested mRNAs are: At_AGO2, AT1G31280; At_AOS, AT5G42650; At_CYP83A1, AT4G13770; At_EDS5, AT4G39030; At_FBA5, AT4G26530; At_GPT2, AT1G61800; At_PAD3, AT3G26830; At_TPS11, AT2G18700.

**Figure S2:**
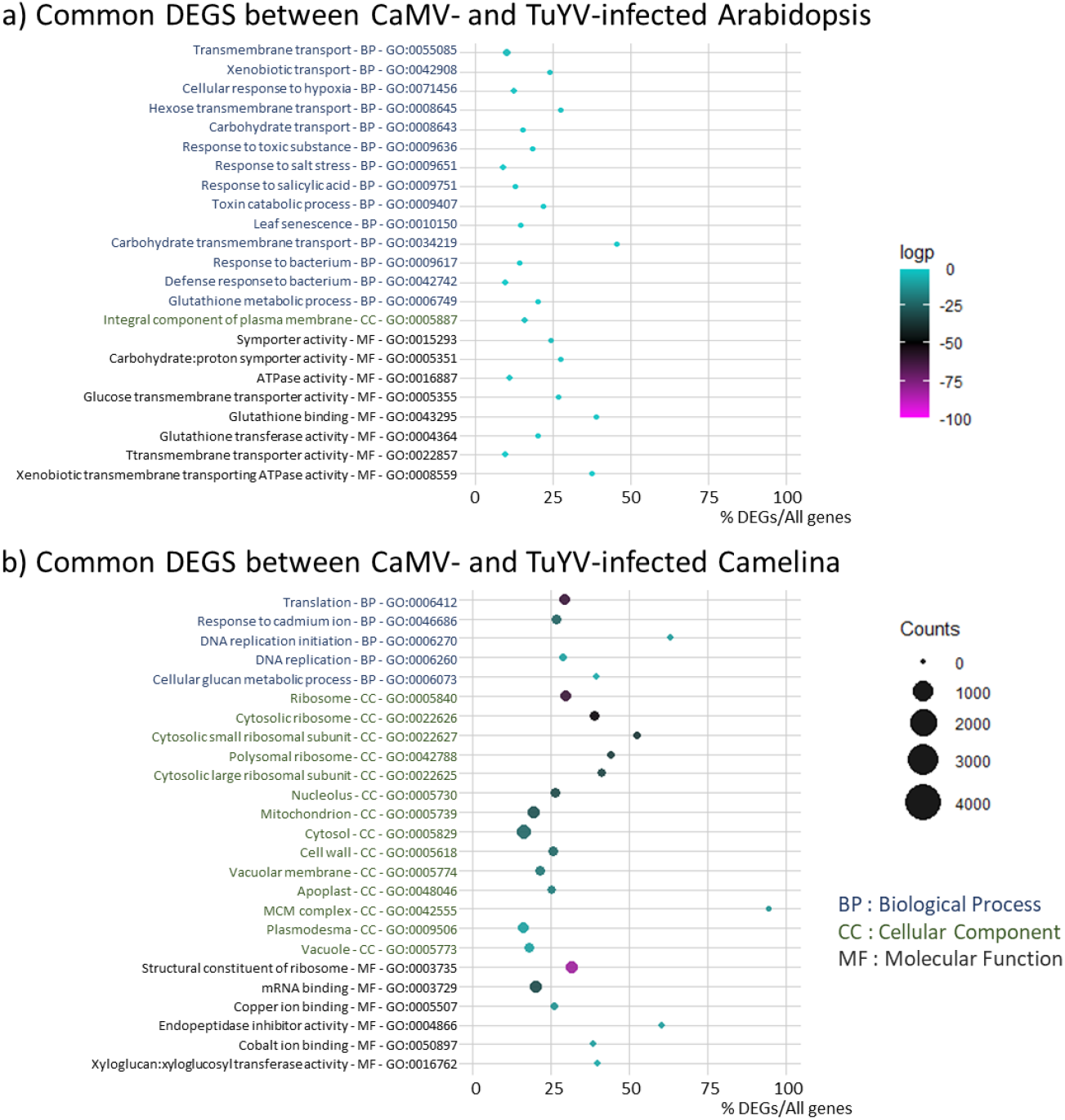
Gene ontology analysis showing the Top 25 GO of deregulated processes. a) common DEGS (n=956) between CaMV-infected and TuYV-infected Arabidopsis and b) common DEGS (n=6,692) between CaMV-infected and TuYV-infected Camelina. GO IDs and corresponding GO terms are specified in the vertical axis. For each category (BP: Biological Process, CC: Cellular Component and MF: Molecular Function), GOs are sorted according to decreasing log2 (1/p-value), also indicated by the color of each spot, in order to place most significantly enriched GOs on top of the graph. The absolute number of DEGs that matched the GO term is indicated by the size of each spot, whereas the horizontal axis shows the ratio of DEGs vs. all genes belonging to the GO term.

### Supplementary heatmaps

Callose deposition is induced via pathogen molecular pattern- and pathogen effector-triggered immunity pathways. We found that expression of genes related to callose deposition was only slightly deregulated in aphid-infested infected plants except for a strong upregulation of *SWEET13* in Camelina infected with CaMV or TuYV, and a slight downregulation of *BHLH89* in CaMV-infected Arabidopsis and Camelina. SWEET13 is a sugar transporter and there is no direct link with callose (at least I did not find any, even if there is callose deposition in the Arabidopsis GO annotation). BHLH89 is a transcription factor that is upstream of callose deposition; (no more information). *UGP1* was virus-specifically upregulated in TuYV-infected Camelina and Arabidopsis and downregulated in CaMV-infected plants. *UGP1* (AT3G03250) is a UDP-glucose pyrophosphorylase and involved in the first synthesis steps of cellulose and other sugar polymers (PMID 29569779) such as callose.

**Figure S3.**
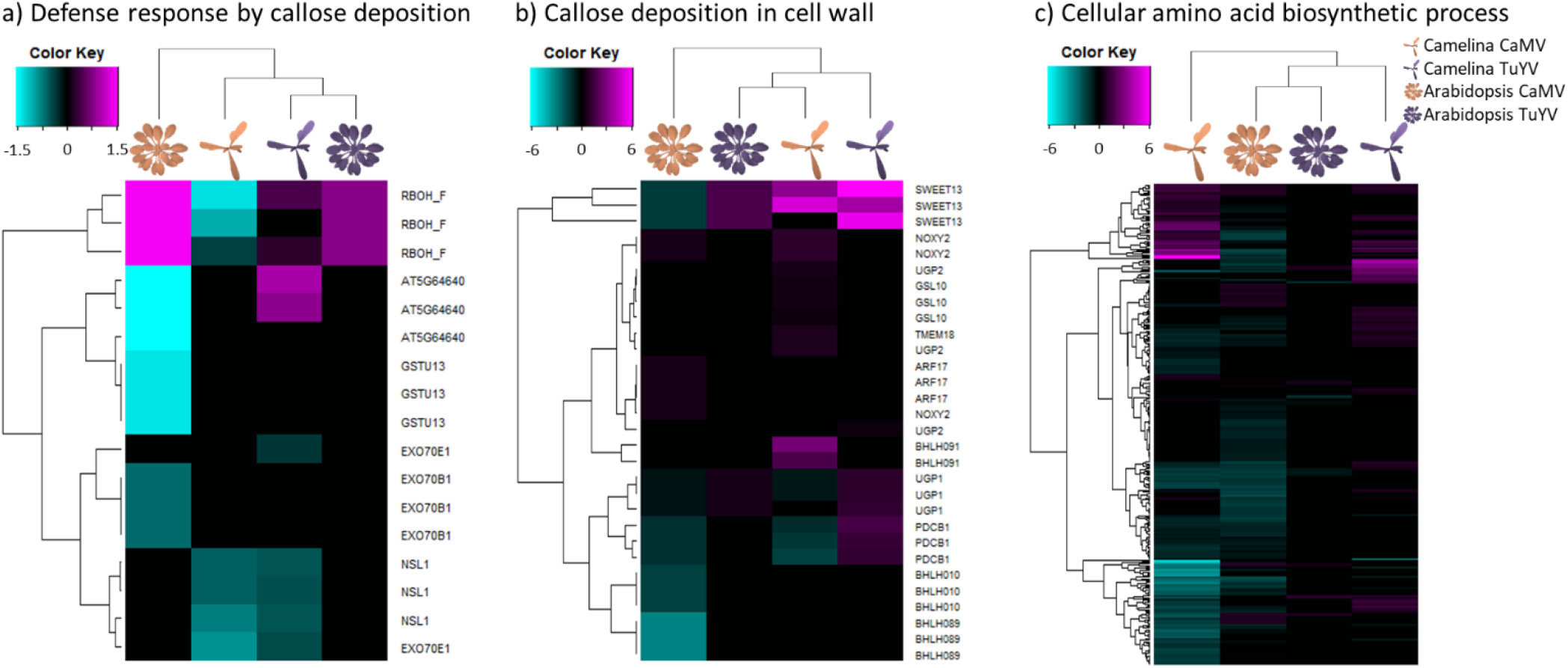
Hierarchical clustering of differentially expressed genes (DEGs) related to a) Defense response by callose deposition (GO:0052542), b) Callose deposition in cell wall (GO:0052543) and c) Cellular amino acid biosynthetic process (GO:0008652) in CaMV- and TuYV-infected *Arabidopsis thaliana* and *Camelina sativa* compared to their mock-inoculated relatives (Supplementary Dataset S2). The color keys show log2fold changes as indicated below the keys in gradients from the minimal value in cyan to the maximal value in magenta.

## Notes

### Competing Interest Statement

The authors have declared no competing interest.

