## Supplementary Data Sets and Sequence Information for "Comparative plant transcriptome profiling of Arabidopsis and Camelina infested with *Myzus persicae* aphids acquiring circulative and non-circulative viruses reveals virus- and plant-specific alterations relevant to aphid feeding behavior and transmission": Supplementary Sequence Information S1 on CaMV and TuYV.docx

**Supplementary sequence information S1.** **Complete genome sequences of CaMV CM1841Rev and TuYV FL1 isolates reconstructed by RNA-seq.** The nucleotides highlighted in yellow represent single nucleotide polymorphism (SNP) positions distinguishing the reconstructed sequences from the reference sequences of CaMV strain CM1841 (V00140) and TuYV isolate BWYV-FL1 (X13063).

>CM1841rev_RNAseq_consensus

GGTATCAGAGCCATGAATCGGTTTAAAAACCAAACTCAAGAGGGTAAAACCTCACCAAAATACGAAAGAGTTCTTAACTCTAAAGATAAAAGATCTTTCAAGATCAAAACTAGTTCCCTCACACCGGTGACCGACAGGTTTACCACCGTAAGGTTTCAGAACAACATCGAATGCGTTTACGCCAACTTCGACTCTCAGCTCAAGTCGTCGTACGATGGTAGATCTAAAAAGATCAAGAATCTAAGCCTTAAAAATCTTAGATGTTATGAAGCCTTCCTCAGGAAGTACCTTCTGGAACAATAAATCTCTCTGAGAATAGTACTCTAACGAGTATCCACAGGAAAAATAATCTTCTGTGTTGAGATGGATTTGTATCCAGAAGAAAATACCCAAAGCGAGCAATCGCAGAATTCTGAAAATAATATGCAAATATTTAAGTCAGAAAATTCGGATGGATTCTCCTCCGATCTAATGATCTCAAACGATCAATTAAAAAATATCTCTAAGACCCAATTAACTTTGGAGAAAGAAAAGATATTTAAAATGCCTAACGTTTTATCTCAAGTTATGAAAAAAGCGTTTAGCAGGAAAAACGAGATTCTTTACTGCGTCTCGACAAAAGAATTATCAGTGGACATTCACGATGCCACAGGTAAGGTATATCTCCCTTTAATCACTAGAGAGGAGATAAATAAAAGACTTTCTAGCTTAAAACCTGAAGTCAGAAAGATCATGTCCATGGTTCATCTTGGAGCGGTCAAAATATTGCTTAAAGCTCAATTTCGAAATGGGATTGATACCCCAATCAAAATTGCTTTAATCGATGATAGAATTAATTCTAGAAGAGATTGCCTTCTCGGTGCAGCCAAAGGTAATCTAGCATACGGTAAGTTTATGTTTACTGTATACCCCAAGTTTGGAATAAGCCTTAATACCCAAAGACTTAACCAAACCTTAAGCCTTATTCATGATTTTGAGAATAAAAATCTTATGAATAAAGGTGATAAAGTTATGACCATAACCTATATCGTAGGATATGCATTAACTAATAGTCATCATAGCATAGATTATCAATCGAATGCTACAATTGAACTAGAAGACGTATTTCAAGAAATTGGAAATGTCCAGCAATCTGACTTTTGTACAATACAAAATGACGAATGCAATTGGGCCATTGATATAGCCCAAAACAAAGCCTTATTAGGAGCTAAAACCCAATCCCAAATTGGTAATAGTCTTCAAATAGGAAACAGTGCTTCATCCTCTAATACTGAAAATGAATTAGCTAGGGTAAGCCAAAACATAGATCTTTTAAAAAACAAATTAAAAGAGATCTGTGGAGAATAAAATGAGCATTACGGGTCAACCGCATGTTTATAAAAAGGATACTATTATTAGACTAAAACCATTGTCTCTTAATAGTAATAATAGAAGTTATGTTTTTAGTTCCTCAAAAGGGAACATTCAAAATATAATTAATCATCTTAACAACCTCAATGAGATTGTAGGAAGAAGCTTACTCGGAATATGGAAGATCAACTCATACTTCGGACTAAGCAAAGACCCTTCGGAGTCCAAATCAAAAAACCCGTCAGTTTTTAATACTGCAAAAAACATTTTTAAAAGTGGGGGGGTTGATTACTCGAGCCAACTAAAGGAAGTAAAATCCCTTTTAGAAGCTCAAAATACTAGAATTAAAAATCTAGAAAATGCAATTCAATCCTTAGATAATAAGATTGAACCAGAGCCCTTAACTAAAGAAGAAGTTAAAGAGCTAAAAGAATCGATTAACTCGATCAAAGAAGGATTAAAGAATATTATTGGCTGAAATGGCTAATCTTAATCAAATCCAGAAAGAAGTCTCTGAAATCCTCAGTGACCAAAAATCCATGAAATCGGATATAAAAGCTATCTTAGAATTGCTAGGATCCCAAAATCCTACTAAAGAAAGCTTAGAAGCCGTTGCAGCGAAAATCGTTAATGACTTAACCAAGCTCATCAATGATTGTCCTTGTAACAAAGAGATATTAGAAGCCTTAGGCAATCAGCCTAAAGAGCAACTAATAGAACAACCTAAAGAAAAAGGCAAAGGCCTTAATCTAGGAAAATATACTTACCCCAATTACGGCGTAGGAAATGAAGAATTAGGATCCTCTGGAAACCCTAAAGCTTTAACTTGGCCTTTCAAAGCTCCAGCAGGATGGCCGAATCAATTTTAGACAGGACCATTAACCGGTTCTGGTATAACCTGGGAGAAGATTGTCTCTCAGAAAGTCAATTTGACCTTATGATAAGGTTAATGGAAGAGTCCCTTGACGGGGACCAAATTATTGATCTAACCTCTCTACCTAGTGATAATTTGCAGGTCGAACAGGTTATGACAACTACCGACGACTCGATCTCGGAAGAATCAGAATTCCTTCTAGCAATAGGAGAAATATCTGAAGACGAAAGTGATTCAGGAGAAGAACCTGAATTCGAACAAGTTCGAATGGATCGAACAGGAGGAACGGAGATTCCCAAAGAAGAAGATGGTGAAGGACCATCTAGATACAATGAGAGAAAGAGAAAGACCCCGGAGGACCGGTACTTTCCAACTCAACCAAAGACCATCCCAGGACAAAAGCAAACGTCTATGGGAATGCTCAACATTGACTGCCAAATCAATCGAAGAACTTTAATCGATGATTGGGCAGCAGAAATCGGATTGATAGTCAAAACCAACAGAGAAGACTATCTTGATCCAGAAACAATACTACTCTTGATGGAACACAAAACATCAGGAATAGCCAAGGAGTTAATCCGAAATACAAGATGGAACCGTACTACCGGCGATATCATAGAACAGGTGATCAATGCAATGTACACCATGTTCTTAGGACTTAACTACTCCGACAACAAGGTTGCTGAAAAGATAGACGAGCAAGAGAAGGCCAAGATCAGAATGACCAAGCTCCAGCTCTTCGACATCTGCTACCTTGAAGAATTTACATGTGATTATGAGAAGAACATGTACAAGACGGAAATGGCGGATTTCCCTGGATACATCAACCAGTACCTGTCAAAAATCCCCATCATAGGAGAAAAAGCGCTAACACGCTTTAGGCATGAAGCCAACGGAACCAGCATCTACAGCTTAGGTTTCGCGGCAAAGATAGTAAAAGAAGAACTATCAAAAATCTGCGACTTATCAAAGAAGCAGAAGAAGTTGAAGAAATTCAACAAGAAATGCTGCAGCATCGGTGAAGCTTCAGTAGAATATGGAGGCAAGAAAACATCCAAGAAGAAGTATCATAAGCGATACAAGAAAAGATATAAGGTCTATAAACCTTATAAGAAGAAGAAGAAATTCCGATCCGGAAAATACTTCAAGCCCAAAGAGAAGAAGGGCTCAAAGCGAAAGTATTGCCCAAAAGGCAAGAAGGACTGCAGATGTTGGATCTGCAATATCGAAGGCCATTACGCCAACGAATGTCCTAATCGACAAAGCTCGGAGAAGGCTCACATCCTTCAACAAGCAGAGAATTTGGGTCTCCAGCCCGTTGAAGAACCCTATGAAGGAGTTCAAGAAGTATTCATCTTAGAATACAAAGAAGAGGAAGAAGAAACCTCTACAGAAGAAAGCGATGATGAATCATCTACTTCTGAAGACTCAGACTCAGATTGAGCAGGTGATGAACGTCACCAATCCCAATTCGATCTACATCAAGGGAAGACTCTACTTCAAAGGATACAAGAAGATAGAGCTTCACTGTTTTGTAGACACGGGAGCAAGCTTATGCATAGCATCCAAGTTCGTCATACCAGAAGAACATTGGGTTAATGCAGAAAGACCAATAATGGTCAAAATAGCAGATGGAAGTTCAATTACCATCAGCAAAGTCTGCAAAGACATAGACTTGATCATAGCCGGCGAGATATTCAAAATTCCCACCGTCTATCAGCAAGAAAGTGGCATCGATTTCATAATCGGCAACAACTTTTGTCAACTGTATGAACCATTCATACAGTTTACAGATAGAGTTATCTTCACAAAGAACAAGTCCTATCCTGTTCATATTACGAAGCTAACAAGAGCAGTGCGAGTAGGCATCGAAGGATTTCTTGAATCAATGAAGAAACGTTCAAAGACTCAGCAACCTGAGCCGGTGAACATTTCGACAAACAAGATAGAAAATCCACTAGAAGAAATTGCTATTCTTTCAGAGGGGAGGAGGTTATCAGAAGAAAAACTTTTCATCACTCAACAAAGAATGCAAAAAATCGAAGAACTACTAGAGAAAGTATGTTCAGAAAATCCATTAGATCCTAACAAGACTAAGCAATGGATGAAAGCTTCAATCAAGCTCAGCGACCCAAGCAAAGCTATCAAGGTTAAACCCATGAAATACAGCCCAATGGATCGTGAAGAATTTGACAAGCAAATCAAAGAGTTACTGGACCTTAAAGTCATTAAACCCAGTAAAAGCCCTCACATGGCACCAGCCTTCTTGGTCAACAATGAAGCCGAGAAGCGAAGAGGAAAGAAACGTATGGTAGTCAACTACAAAGCTATGAACAAAGCCACCATAGGAGACGCATACAATCTTCCCAACAAAGACGAGTTACTTACACTCATTCGAGGAAAGAAGATCTTTTCTTCCTTCGACTGTAAGTCCGGATTCTGGCAAGTTCTACTTGATCAAGAATCAAGACCTCTAACGGCATTCACATGTCCACAAGGTCACTACGAATGGAATGTGGTCCCTTTCGGCCTAAAGCAGGCACCATCCATATTCCAGAGACACATGGACGAAGCATTTCGTGTGTTCAGAAAATTCTGTTGCGTGTATGTCGACGACATCCTCGTATTCAGTAACAACGAAGAAGATCACCTACTTCACGTAGCAATGATCTTACAAAAGTGCAATCAACATGGAATCATTCTTTCCAAGAAGAAAGCACAACTCTTCAAGAAGAAGATAAACTTCCTTGGTCTAGAAATAGATGAAGGAACACACAAGCCTCAAGGACATATCTTGGAACATATCAACAAATTCCCAGATACCCTTGAAGACAAGAAGCAACTTCAGAGATTCTTAGGCATCCTAACATATGCCTCTGATTATATCCCGAAGCTAGCTCAAATCAGAAAGCCTCTGCAAGCCAAGCTTAAAGAAAATGTTCCATGGAAATGGACAAAGGAGGACACCCTCTACATGCAAAAGGTGAAGAAAAATCTGCAAGGATTTCCTCCACTACATCATCCCTTACCAGAGGAAAAGCTGATCATCGAGACCGACGCATCAGACGACTACTGGGGAGGTATGTTAAAAGCTATCAAAATTAACGAAGGTACTAATACCGAGTTAATTTGCAGATACGCATCTGGAAGCTTTAAAGCTGCAGAAAGGAATTACCACAGCAATGACAAAGAGACATTGGCGGTAATAAATACTATAAAGAAATTCAGTATTTATCTAACTCCTGTTCATTTTCTGATTAGGACAGATAATACTCATTTCAAGAGTTTTGTTAACCTTAATTACAAAGGAGATTCAAAACTTGGAAGAAACATCAGATGGCAAGCATGGCTTAGCCACTATTCGTTTGATGTTGAACATATTAAAGGAACCGACAACCACTTTGCGGACTTCCTTTCAAGAGAATTCAATAAGGTTAATTCCTAATTGAAATCCGAAGATAAGATTCCCACACACTTGTGGCTGATATCAAAAAGGCTACTACCTATATAAACACATCTCTGGAGACTGAGAAAATCAGACCTCCAAGCATGGAGAACATAGAAAAACTCCTCATGCAAGAGAAAATACTAATGCTAGAGCTCGATCTAGTAAGAGCAAAAATAAGCTTAGCAAGAGCTAACGGCTCTTCGCAACAAGGAGACCTCCCTCTCCACCGTGAAACACCGGTAAAAGAAGAAGCAGTTCATTCTGCACTGGCCACTTTTACGCCAACTCAAGTAAAGGCTATTCCAGAGCAAACGGCTCCTGGTAAAGAATCAACAAATCCGTTGATGGCTAGTATCTTGCCAAAAGATATGAACCCAGTTCAAACTGGGATAAGGCTTGCAGTGCCAGGGGACTTTTTACGTCCTCATCAGGGAATTCCAATCCCACAAAAATCTGAGCTTAGCAGCATAGTTGCTCCTCTCAGAGCAGAATCGGGTATTCAACACCCTCATATCAACTACTACGTTGTGTATAACGGTCCACACGCCGGTATATACGATGACTGGGGTTGTACAAAGGCGGCAACAAACGGCGTTCCCGGAGTTGCACACAAGAAGTTTGCCACTATTACAGAGGCAAGAGCAGCAGCTGACGCGTACACAACAAGTCAGCAAACAGACAGGTTGAACTTCATCCCCAAAGGAGAAGCTCAACTCAAGCCCAAGAGCTTTGCGAAGGCCTTAACCAGCCCACCAAAGCAAAAAGCCCACTGGCTCACGCTAGGAACCAAAAGGCCCAGCAGTGATCCAGCCCCAAAAGAGATCTCCTTTGCCCCGGAGATCACCATGGACGACTTTCTCTATCTCTACGATCTAGGAAGAAAGTTCGACGGAGAAGGTGACGATACCATGTTCACCACCGATAATGAGAAGATTAGCCTCTTCAATTTCAGAAAGAATGCTGACCCACAGATGGTTAGAGAGGCCTACGCGGCAGGTCTCATCAAGACGATCTACCCGAGTAATAATCTCCAGGAGATCAAATACCTTCCCAAGAAGGTTAAAGATGCAGTCAAAAGATTCAGGACTAACTGCATCAAGAACACAGAGAAAGATATATTTCTCAAGATCAGAAGTACTATTCCAGTATGGACGATTCAAGGCTTGCTTCATAAACCAAGGCAAGTAATAGAGATTGGAGTCTCTAAGAAAGTAGTTCCTACTGAATCAAAGGCCATGGAGTCAAAAATTCAGATCGAGGATCTAACAGAACTCGCCGTGAAGACTGGCGAACAGTTCATACAGAGTCTTTTACGACTCAATGACAAGAAGAAAATCTTCGTCAACATGGTGGAGCACGACACTCTCGTCTACTCCAAGAATATCAAAGATACAGTCTCAGAAGACCAAAGGGCTATTGAGACTTTTCAACAAAGGGTAATATCGGGAAACCTCCTCGGATTCCATTGCCCAGCTATCTGTCACTTCATCAAAAGGACAGTAGAAAAGGAAGGTGGCACCTACAAATGCCATCATTGCGATAAAGGAAAGGCTATCGTTCAAGATGCCTCTGCCGACAGTGGTCCCAAAGATGGACCCCCACCCACGAGGAGCATCGTGGAAAAAGAAGACGTTCCAACCACGTCTTCAAAGCAAGTGGATTGATGTGATATCTCCACTGACGTAAGGGATGACGCACAATCCCACTATCCTTCGCAAGACCCTTCCTCTATATAAGGAAGTTCATTTCATTTGGAGAGGACACGCTGAAATCACCAGTCTCTCTCTACAAATCTATCTCTCTCTATTTTCTCCATAATAATGTGTGAGTAGTTCCCAGATAAGGGAATTAGGGTTCTTATAGGGTTTCGCTCACGTGTTGAGCATATAAGAAACCCTTAGTATGTATTTGTATTTGTAAAATACTTCTATCAATAAAATTTCTAATTCCTAAAACCAAAATCCAGTACTAAAATCCAGATCACCTAAAGTCCCTATAGATCTTTGTCGTGAATATAAACCAGACATGAGACGACTAAACCTGGAGCCCAGACGCCGTTCGAAGCTAGAAGTACCGCTTAGGCAGGAGGCCGTTAGGGAAAAGATGCTAAGGCAGGGTTGGTTACGTTGACTCCCCCGTAGGTTTGGTTTAAATATGATAAAGTGGACGGAAGGAAGGAGGAAGACAAGGAAGGATAAGGTTGCAGGCCCTGTGCAAGGTAAGAAGATGGAAATTTGATAGAGGTACGTTACTATACCTATACTATACGCTAAGGGATGCTTGTATTTTACCCTATACCCCCTAATAACCCCTTATCGATTTTAAGAAATAATCCGCATAAGCCCCCGCTTAAAAAATT

>TuYV_RNAseq_consensus

ACAAAAGAAACCAGGAGGGAATCCTTAGTTGATGCAATTTCTCGCTCACGATAACTTTCACACTTTGCAAGTCAAGAAAGTCCGATTCCTCCACCCTCAACAAGAAGTGTTTCTTTTAGCAGGTTTATTGCTCAATATAAAACAATTcGTACGAGCAATCAAAGAGCGCAATAATGTATTCAAAATTGATGTTTTTCTTCGCTCTTTGCTCTATCAGCTTCCTTTTCACCTCGGAAGCTGCTTCCACGATGCTCCTCGAGAGCTCATACCTGCCACTGAACCAGAGTTATGCGCCTGGTTTTCTTTACAAACGGGATATGCTCCCGCCTCCACTTCAGGCCGTGTTAACTTACACGTGCCCGGAACCAAGACCTCTCGCAGAAGAATCATACAACGATCTTTTGCGAGCGATTTCTCAGAAAAGCTCAAGCGATTTCCAGAATGCTTATTCGTTAGCCTTGAGCTTTTCCAGCGACTTCTATCAACATGGACTAAAGACGTTGAAAGACGTATCTTTTTTAGCTGTCGAGAAATTCCTCTGGGGTCTGACACGCTTATGGAGCTCGCTAATCTTGGCGAGTTTCTCCGCGTTATGGTGGTTGGTGAGCAATTTCACAACTCCCGTCTTCTGTCTCGCCTTGCTGTACACTGTTACAAAATATATGGTGAAGACGGTTTCATTTCTTTTTGGAGGATTGCCAATCTGGATCATTTCGATTGCTTTCTCACTCCTGAAGAAATCCTTTTCAGCTCTTCGGTCTACACCGAAATGTTTGTATGAAAAGGCCATAGACGGTTTCAAGAGTTTCACTATCCCGCAGAGTCCCCCAAAATCTTGCGTGATTCCCATCACCCACGCAAGCGGAAACCACGCTGGCTATGCCAGTTGTATCAAGCTATATAACGGAGAAAATGCTCTAATGACGGCAACTCACGTCCTACGTGATTGCCCCAACGCCGTGGCTGTTTCCGCCAAAGGACTCAAAACGCGGATTCCACTCGCAGAATTCAAAACAATCGCGAAATCCGACAAAGGTGATGTTACCCTCCTTCGCGGCCCCCCCAATTGGGAAGGACTGTTGGGCTGTAAAGCGGCCAACGTTATAACAGCTGCcAACCTAGCGAAATGCAAAGCATCCATATACTCTTTTGACAGAGATGGCTGGGTTAGCAGTTATGCCGAGATCGTgGGCTCAGAAGGTACAGATGTTATGGTTCTGAGCCATACGGAAGGAGGACACTCCGGAAGCCCCTACTTCAATGGTAAAACCATCTTGGGGGTTCATTCAGGTGCCAGTGCTACTGGAAATTACAATTTAATGGCACCAATCCCATCCCTCCCCGGACTTACTAGTCCGACTTATGTGTTCGAAACCACCGCACCACAAGGAAGAGTTTTCGCACAAGAAGATATCGCTGAAATCGAAGGCCTCTATGCACAAGTAATGAAAAGAGTTCAACAAGCGGAAGATTTCAAACCCAAAACTGGAAAGTATTGGGGTGATATGGAGGATGATGAAGACATTTTCTTCGAAAGCAAAGAAGATCTGTCGGGAAACGGAGTGCGCGGCACCGTCCGCGGAACAAACGGAGAAGGCAGCTCCACCCCAAAGACAAGCAACGTCGATGGGAAAGAGATGATGGAGAAAATAATCTCATCTCTAGTGGGAAAGATAAATCTCGAGAACATCGAGAGGAAAGTGATAGAGGAGATCTCCGCGAAAGCGATGAAAACTCCGAAATCCCGCCGCAGAAGAGCCCCAAAGAAACAGCCGGAGAGTTCGAAAGATACTTCTCCTCGCTCTACAACTGGGAAGTACCAACCTCCCCACGTGAGGTCCCCGGCTTCCGTCACTGCGGCAAACTGCCCCAATACTACCACCCCAAGCAAAAAGAAGAATCTAGCTGGGGGAAGACCCTCGTCGGGAACCATCCCGCGTTGGGTGAGAAAACAAGCGGCTTCGGCTGGCCCAAGTTCGGCCCCGAAGCAGAACTGAAGAGCCTGCGACTGCAGGCTTCACGGTGGCTGGAACGCGCCCAGTCCGCAGAAATACCCTCTGACGCTGAGAGGGAGCGTGTGATTCAAAAGACCGCAGATGTTTACCATCCTTGCCAAACAAATGGGCCTGCGGCAACCCGAGGAGGAACACTAACCTGGAACAACTTCATGATTGATTTCAAACAGGCAGTGTTCTCGCTGGAGTTCGATGCCGGAATCGgcgTCCCCTATATTGCCTATGGCAAGCCCACACACCGTGGGTGGGTTGAAGACCAGAAACTCCTTCCAATCCTAGCTCAATTGACCTTCTTCCGACTACAGAAGATGTTGGAGGTCAATTTCGAAGATATGGGACCTGAGGAGCTGGTCCGGAACGGTTTGTGTGATCCCATCCGATTATTCGTGAAGGGTGAGCCGCACAAGCAAGCGAAGCTCGATGAAGGCCGCTACCGCCTCATAATGAGCGTTTCCCTCGTGGATCAACTGGTAGCCCGGGTTCTGTTTCAAAATCAGAACAAGCGGGAAATTGCCCTGTGGAGGGCCATCCCCAGCAAACCCGGTTTTGGCTTGTCTACGGATGAGCAAGTGCTGGACTTTGTGGAAAGTCTGGCCCGTCAAGTAGGCACCACTACGACAGAGGTGGTTGCCAATTGGAAcAATTACTTGACGCCCACGGATTGCTCCGGTTTTGACTGGAGTGTTGCGGATTGGATGCTTCACGACGACATGATCGTCCGCAACAGACTTACCATCGACCTCAACCCCGCTACAGAAAGATTAAGATCTTGCTGGTTGAGGTGCATTTCAAACTCAGTATTGTGCCTGAGTGATGGCACCCTTTTAGCCCAAATTCATCCGGGCGTTCAGAAGAGTGGGAGCTATAATACATCAAGCTCCAACTCCCGGATCCGAGTTATGGCCGCCTTCCACACAGGTGCCATCTGGGCTATGGCGATGGGTGATGATGCCCTCGAGTCCAATCCCGCTGACCTAGCAGCGTACAAGAAACTAGGTTTCAAGGTTGAGGTTTCCGGACAACTGGAATTCTGCTCTCAcATCTTTAGAGCGCCGGACCTCGCCCTCCCTGTGAACGAAAATAAGATGATCTACAAACTGATCTATGGCTATAATCCAGGGAGCGGAAACGCTGAGGTAGTTTCAAACTACTTGGCCGCTTGTTTCTCAGTTCTGAACGAGTTGCGGCATGATCCAGCGTCCGTTGAACTTCTTTACTCGTGGTTAGTCGATCCGGTGCTACCACAAAAGATACCAGGAGAGTAAAGAAGAAGAAAGTCAGCTTACATTGAAATTTTAAAGAGGTTTCTGCAACAGTAAGAGACTTAAGCAAACCCAATTAAAGATACAACGGATTACAAATTCCTAGCAGGCTTCGCCGCAGGCTTCGTTTCATCGATACCAATATCCGTGATCAGTATCTATTTCATCTACCTAAGAATCTCCAAACACGTACGCGAAATCGTTAATGAATACGGTCGTGGGTAGGAGAATTATCAATGGAAGAAGACGACCACGCAGGCAAACACGACGCGCTCAGCGCCCTCAGCCAGTGGTTGTGGTCCAAACCTCTCGGGCAACACAACGCCGACCTAGACGACGACGAAGAGGTAACAACCGGACAGGAAGAACTGTTCCTACCAGAGGAGCAGGTTCGAGCGAGACATTTGTTTTCTCAAAAGACAATCTCGCGGGAAGTTCCAGCGGAGCAATCACGTTCGGGCCGAGTCTATCAGACTGCCCGGCATTCTCTAATGGAATGCTCAAGGCCTACCATGAGTATAAAATCTCAATGGTCATTTTGGAGTTCGTCTCCGAAGCCTCTTCCCAAAATTCCGGTTCCATCGCTTACGAGCTGGACCCACACTGTAAACTCAACTCCCTTTCCTCAACTATCAACAAGTTCGGGATCACAAAGCCCGGGAAAAGGGCGTTTACAGCGTCTTACATCAACGGAACGGAATGGCACGACGTTGCCGAGGACCAATTCAGGATCCTCTACAAAGGCAATGGTTCTTCATCGATAGCTGGTTCTTTCAGAATCACCATTAAGTGTCAATTCCACAACCCCAAATAGGTAGACGAGGAACCCGGCCCTAGCCCAGGGCCTTCTCCCTCTCCACAACCCACACCCCAAAAGAAATATCGTTTTATCGTCTATACTGGAGTCCCCGTGACTCGTATAATGGCTCAATCTACGGATGATGCCATCTCTTTGTATGATATGCCGTCCCAACGGTTTCGCTACATAGAGGACGAGAACATGAACTGGACGAACCTCGATTCTCGATGGTATTCCCAGAATTCTTTGAAAGCCATCCCGATGATAATAGTGCCAGTCCCTCAAGGTGAGTGGACCGTGGAAATATCGATGGAGGGGTATCAACCAACCTCAAGCACCACAGATCCTAACAAGGACAAACAAGATGGTCTCATCGCCTACAACGATGATCTTAGTGAAGGTTGGAACGTGGGGATTTACAATAATGTGGAGATAACCAACAACAAGGCCGATAATACTTTGAAGTATGGCCACCCAGACATGGAACTCAATGGCTGTCATTTCAATCAAGGACAGTGTCTGGAAAGAGATGGAGATTTGACTTGTCATATCAAGACGACTGGTGACAATGCCTCCTTCTTTGTTGTTGGACCCGCTGTCCAGAAGCAATCTAAATATAATTACGCCGTTTCGTACGGAGCCTGGACAGATCGGATGATGGAGATAGGGATGATCGCCATAGCACTTGATGAACAAGGCTCATCCGGTTCCGTAAAGACAGAAAGACCAAAGAGAGTTGGGCACTCCATGGCAGTCTCAACCTGGGAGACTATAAAATTGCCGGAGAAGGGAAACTCCGAGGGATACGAAACCAGTCAAAGACAAGACTCTAAAACTCCTCCCACAGCTAGTGGGGGTTCTGACACGCTGGACGTCGAAGAAGGAGGCTTGCCCCTTCCTGTTGAAGAAGAGATCCCCGATTTTGTTGGGGATAACCCCTGGTCTGACTTATCGACTAAGAATTCACAGGAAGAAGAGGCTATGTCATCAGAGAGTGGTCTTAGACCCCAGTTGAAGCCTCCTGGTCTGCCAAAACCTCAACCGATCAGAACGATTCGAAACTTCGATCCAACACCGGATTTGGTTGAAGCGTGGCGACCCGATGTGAACCCCGGATATTCCAAAGCAGATGTGGCAGCCGCTACTATCATCGCCGGGGGTTCCATCAAAGACGGCCGTTCTATGATTGATAAACGAAATAAAGCTGTGTTAGACGGTCGCAAGAGTTGGGGTTCCTCCTTGGCTTCCTCCCTCACGGGTGGTACGCTCAAGGCCTCCGCCAAGTCGGAGAAGCTTGCCAAACTTACCACGAGTGAAAGGGCAAGGTATGAACGGATTAAGCGTCAGCAAGGCTCCACAAGAGCCTCGGAATTCCTAGAATCACTTCTAGCTGGCGAAGACCCCGACTCAAGGTTCTGAAGGGATACAACCTGACCCTTCCCGGTCCAGATGAACCCGTCCAAATCATCATCGTCAAGCCAGGGACTTTAAACTGGAACGAATCCGTTTTACGGATAGGCAACGAGTGTTTTACGCTGGGAGAAATCCCTACGGCACTTCGGTGT
